# Early post-zygotic mutations contribute to congenital heart disease

**DOI:** 10.1101/733105

**Authors:** Alexander Hsieh, Sarah U. Morton, Jon A.L. Willcox, Joshua M. Gorham, Angela C. Tai, Hongjian Qi, Steven DePalma, David McKean, Emily Griffin, Kathryn B. Manheimer, Daniel Bernstein, Richard W. Kim, Jane W. Newburger, George A. Porter, Deepak Srivastava, Martin Tristani-Firouzi, Martina Brueckner, Richard P. Lifton, Elizabeth Goldmuntz, Bruce D. Gelb, Wendy K. Chung, Christine E. Seidman, J. G. Seidman, Yufeng Shen

**Author notes:** Corresponding author. (Y.S.). Equal contribution (Alexander Hsieh, Sarah U. Morton, Jon A.L. Willcox, Wendy K. Chung, Christine E. Seidman, J. G. Seidman, Yufeng Shen). (A.H.), (S.U.M.), (J.A.L.W.), (J.M.G.), (A.C.T.), (H.Q.), (S.D.), (D.M.), (E.G.), (K.B.M.), (D.B.), (R.W.K.), (J.W.N.), (G.A.P.), (D.S.), (M.T.F.), (M.B.), (R.P.L.) (E.G.) (B.D.G.) (W.K.C.) (C.E.S.), (J.G.S.), (Y.S.).

## Abstract

**Background:** The contribution of somatic mosaicism, or genetic mutations arising after oocyte fertilization, to congenital heart disease (CHD) is not well understood. Further, the relationship between mosaicism in blood and cardiovascular tissue has not been determined.

**Results:** We developed a computational method, Expectation-Maximization-based detection of Mosaicism (EM-mosaic), to analyze mosaicism in exome sequences of 2530 CHD proband-parent trios. EM-mosaic detected 326 mosaic mutations in blood and/or cardiac tissue DNA. Of the 309 detected in blood DNA, 85/97 (88%) tested were independently confirmed, while 7/17 (41%) candidates of 17 detected in cardiac tissue were confirmed. MosaicHunter detected an additional 64 mosaics, of which 23/46 (50%) among 58 candidates from blood and 4/6 (67%) of 6 candidates from cardiac tissue confirmed. Twenty-five mosaic variants altered CHD-risk genes, affecting 1% of our cohort. Of these 25, 22/22 candidates tested were confirmed. Variants predicted as damaging had higher variant allele fraction than benign variants, suggesting a role in CHD. The frequency of mosaic variants above 10% mosaicism was 0.13/person in blood and 0.14/person in cardiac tissue. Analysis of 66 individuals with matched cardiac tissue available revealed both tissue-specific and shared mosaicism, with shared mosaics generally having higher allele fraction.

**Conclusions:** We estimate that ~1% of CHD probands have a mosaic variant detectable in blood that could contribute to cardiac malformations, particularly those damaging variants expressed at higher allele fraction compared to benign variants. Although blood is a readily-available DNA source, cardiac tissues analyzed contributed ~5% of somatic mosaic variants identified, indicating the value of tissue mosaicism analyses.

## Background

Mosaicism results from somatic mutations that arise post-zygotically in an early embryonic cell, resulting in two or more cell populations with distinct genotypes in the developing embryo {Biesecker 2013}. The developmental status of the early embryonic cell at the time of mutagenesis determines the proportion of variant-carrying cells and the tissue distribution of these cells in the post-natal child {Acuna-Hidalgo 2015}. While germline variants have a variant allele frequency (VAF) of 0.5, somatic mosaic variants have a significantly lower VAF.

Post-zygotic mosaic mutations have been implicated in several diseases including non-malignant developmental disorders such as overgrowth syndromes {Poduri 2013; Lindhurst 2012; Kurek 2016}, structural brain malformations {Poduri 2012; Jamuar 2014; Riviere 2012; Lee 2012}, epilepsy {Stosser 2018}, and autism spectrum disorder {Lim 2017; Krupp 2017; Freed 2016; Dou 2017}. Recent analyses also identified mosaic variants in a cohort of patients with congenital heart disease (CHD) {Manheimer 2018}, but the prevalence of these was far less than germline variants (CHD) {Zaidi 2013; Homsy 2015; Jin 2017; Zaidi 2017}.

Assessment of the frequency of mosaicism in human disease is confounded by technical issues, including differences in sequencing depth, DNA sources, and variant assessment pipelines. Low levels of mosaicism can escape the detection threshold of traditional sequencing methods with standard read depths, while post-zygotic mutations with a higher percentage of affected cells are difficult to discriminate from germline *de novo* mutations {Acuna-Hidalgo 2015}. All of these issues can lead to substantially different conclusions. For example, analyses of mosaicism in autism spectrum disorder was recently assessed from whole exome sequence (WES) data from whole blood DNA from 2506 families (proband, parents and unaffected sibling; trios and quads) in the Simons Simplex Collection (SSC) {Fischbach 2010}. The primary sequence data were analyzed by three groups; one that identified a protein-coding somatic mosaic variant rate of 0.074 per individual {Freed 2016}, another that found a mosaic rate of 0.059 per individual {Lim 2017}, and a third group that reported a mosaic rate of 0.125 per individual {Krupp 2017}. This disparity suggests the need for more systematic mosaic mutation detection methods that account for dataset-specific confounding factors.

By contrast, analyses of affected tissues can improve the sensitivity and specificity of detection of somatic mosaicism. In cancer, methods to detect these events, such as MuTect {Cibulskis 2013}, compare tumor and benign tissues from the same patient. Mosaicism has also been demonstrated from the analyses of unpaired samples with cancer and other pathologies {Sun 2018; Huang 2017; Smith 2015} by the demonstration of variants in affected tissues that are absent from blood-derived DNA {Symoens 2017; McDonald 2018}. With access to cardiac tissues from patients with CHD obtained during surgical repair, we hypothesized that analyses of mosaicism in cardiac tissue might improve insights into the causes of this common congenital anomaly. As many cardiomyocyte lineages share a mesodermic origin with blood cells but exit the cell cycle during embryogenesis, we also sought to determine if mosaicism in the heart exhibited distinct patterns of mosaicism with regard to variant frequency and allele fractions.

In this study, we developed a computational method, EM-mosaic (Expectation-Maximization-based detection of Mosaicism), to detect mosaic single nucleotide variants (SNVs) using WES of proband and parent DNA. To optimize this method, we measured mosaic detection power as a function of sequencing depth. We compared EM-mosaic and MosaicHunter {Huang 2014} to investigate mosaicism in 2530 CHD proband-parent trios from the Pediatric Cardiac Genomics Consortium (PCGC) {Jin et al 2017}, using exome sequences derived from blood-derived DNA. We detected predicted deleterious mosaic mutations in genes involved in known biological processes relevant to CHD or developmental disorders in 1% of probands. The accuracy of these mosaic variant detection algorithms was assessed using an independent re-sequencing method. We found that among high-confidence mosaic mutations in CHD-relevant genes, likely-damaging variants tended to have higher VAF than likely-benign variants.

In parallel we assessed mosaicism by EM-mosaic and MosaicHunter in 70 discarded tissues from several heart regions obtained from 66 probands who underwent cardiac surgical repairs. While VAF varied significantly (>3 fold) between blood and cardiovascular tissue at about 60% of sites, in general mosaic variants with high (>15%) VAF were more likely shared between blood and cardiac tissue than variants with lower VAF.

## Results

### High-accuracy detection of mosaic mutations in WES data using EM-mosaic

We analyzed whole exome sequence (WES) data from 2530 CHD proband-parent trios {Homsy 2015; Jin 2017} (**Table S1**). Among this cohort, 1205 probands had CHD with neurodevelopmental disorders (NDD) and/or extracardiac manifestations (EM), 788 had isolated CHD at the time of enrollment, 539 had undetermined NDD status due to young neonatal age at the time of enrollment, and 9 subjects had incomplete data (**Table S2**).

Previous WES analyses {Jin et al 2017} identified 1742 germline *de novo* SNVs among 838 cases with NDD and/or EM, 516 isolated cases, 644 cases of unknown NDD status, and 7 with incomplete data. These *de novo* variants were identified using the Genome Analysis Toolkit (GATK) pipeline {McKenna 2010; DePristo 2011} assuming a germline diploid model in which the expected VAF is 0.5. This model has limited sensitivity to detect mosaic mutations for which the fraction of alternative allele reads is significantly below 0.5, especially because *de novo* variants with VAF<0.2 were excluded to reduce false discovery.

To efficiently capture mosaic variants with VAF<0.4, we developed a new method (EM-mosaic) to detect mosaic variants in WES sequence of a proband and parents (trios). Potential mosaic variants were identified in WES sequence data using SAMtools *mpileup* {Li 2009} with settings designed to capture sites with VAF between 0.1-0.4 and merged with the variants found by the GATK pipeline {Jin et al 2017} (**Fig S1**) to create a union variant set. To reduce the elevated false positive rate inherent in low-VAF calls, we applied a set of empirical filters to remove likely technical artifacts due to sequencing errors associated with repetitive and/or low complexity sequences. We then manually inspected *de novo* SNVs with VAF<0.3 (n=582) using IGV and filtered out an additional 188 likely false positives. After preprocessing and outlier removal, the remaining 2971 *de novo* SNVs were used as input to our mosaic detection model.

Among the 2971 *de novo* SNVs, this pipeline identified 309 sites as candidate mosaics based on posterior odds score (**Fig 1A-B, Table S3**), including 50 sites that were previously reported as germline *de novo* variants {Jin et al 2017}. An additional 86 sites were identified as having posterior odds below our threshold of 10 but greater than 1 (**Fig S2A, S2B**), including a *ZEB2* variant with posterior odds 4.7 that was previously confirmed via ddPCR {Manheimer 2018}. Among these 86 variants, 53 are likely mosaic and 33 are likely germline (**Fig S2B**). We chose not to include these sites since there was insufficient evidence to confidently resolve them individually as mosaic or germline.

**Fig 1.**
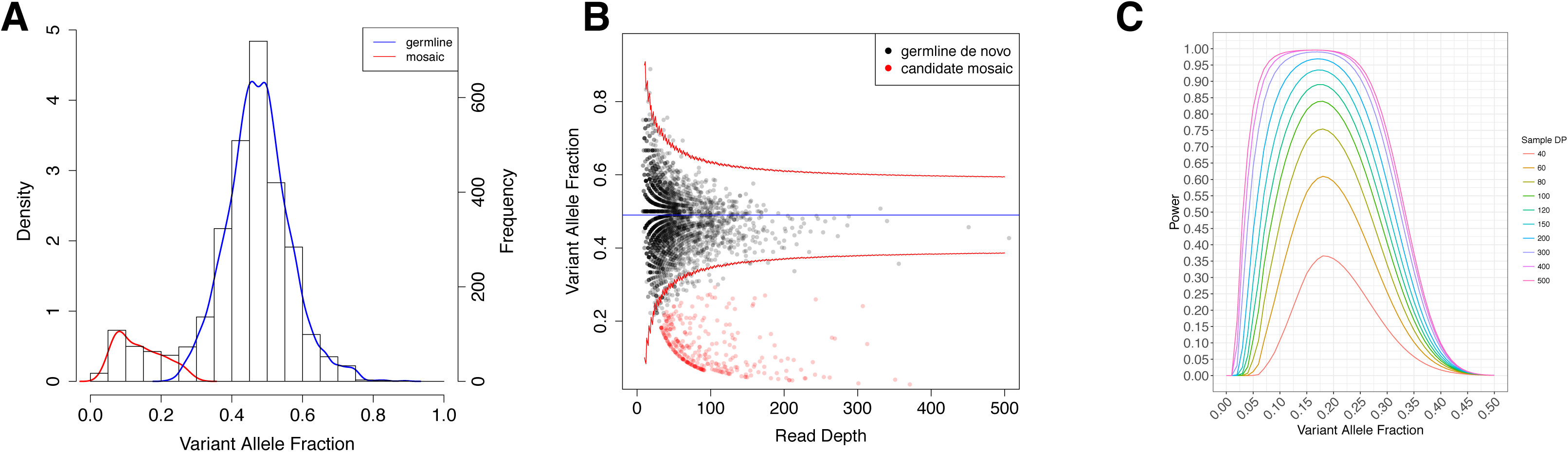
Mosaic detection by Expectation-Maximization. **(A)** Expectation-Maximization (EM) Estimation to decompose the variant allele fraction (VAF) distribution of our input variants into mosaic and germline distributions. The EM-estimated prior mosaic fraction was 12.15% and the mean of the mosaic VAF distribution was 0.15. (**B**) Read depth vs. VAF distrubution of individual variants. The blue line denotes mean VAF (0.49) and the red lines denote the 95% confidence interval under our Beta-Binomial model. Mosaic variants are defined as sites with posterior odds > 10, corresponding to a False Discovery Rate of 9.1%.Germline variants are represented in black and mosaic variants are represented in red. (**C**) Estimated mosaic detection power as a function of average sample depth for values between 40x and 500x.

### Mosaic mutations found in blood derived DNA with MosaicHunter

We also employed MosaicHunter, which uses a Bayesian genotyping algorithm with a series of stringent filters (see Supplemental Methods) for discovering mosaic variants using WGS genotype information from trios. {Huang 2017} Among the 2530 CHD trios, MosaicHunter identified an initial set of 58976 sites showing evidence of mosaicism, including 214 high-confidence variants located in coding regions. (**Fig S3**). After applying a minimum likelihood ratio (LR) cutoff of 80 for distinguishing mosaic from germline mutation, and additional heuristic filters (**Supplemental Methods**), MosaicHunter identified 116 coding sites (**Table S4**) or 0.05 mosaics /individual.

Of the mosaic candidates detected by MosaicHunter, 58/116 (50%) were also identified by EM-mosaic while 58/116 (50%) candidates were unique to MosaicHunter (**Table 1; Fig S4**). Of the 58 candidates unique to MosaicHunter, 35 were filtered out by EM-mosaic on the basis of insufficient alternate allele read support, 16 had a non-zero allelic depth in the parents, and 7 failed quality filters. The 251 candidates unique to EM-mosaic were discarded by the MosaicHunter pipeline during BAM reprocessing (n=13), quality filtering (n=146), application of LR cutoff (22), or were not called due to inadequate read depth (n=70) (**Fig S3**).

**Table 1:** Mosaic variant detection by EM-Mosaic, MosaicHunter and validated by PCR product sequencing

|  |  | Union | Shared | Unique |  |
| --- | --- | --- | --- | --- | --- |
|  |  |  |  | EM-Mosaic | MosaicHunter |
| Mosaic Variants (total)* |  | 315 (332) | 56 (57) | 218 (240) | 29 (35) |
| Mosaic Candidates |  | 367 | 58 | 251 | 58 |
| Mosaic Candidate VAF mean (SD) |  | 0.13 (0.06) | 0.12 (0.05) | 0.13 (0.06) | 0.10 (0.05) |
| MiSeq Confirmation | Total Tested | 143 | 22 | 75 | 46 |
|  | Mosaic | 108 | 21 | 64 | 23 |
|  | Germline | 3 | 0 | 3 | 0 |
|  | No Variant | 32 | 1 | 8 | 23 |
| Validation Rate |  | 76% | 95% | 85% | 50% |

\*Estimated number of mosaic variants found among 2530 CHD probands (total number of mosaic variants detected by EM-mosaic and MosaicHunter).

### Sequence confirmation of candidate mosaic variants and estimation of mosaicism in CHD

From the 367 high-confidence EM-mosaic and/or MosaicHunter SNVs, we selected 143 candidates (97 identified by EM-mosaic; 68 identified by MosaicHunter) for experimental confirmation using MiSeq amplicon resequencing (**Table S5; Table S11 and S12; Methods**). DNA fragments encompassing the putative mosaic variant were PCR-amplified from proband and each parent DNA, sequenced on an Illumina MiSeq next generation sequencer and VAF was calculated for each individual. These candidate mosaics included SNVs on the extremes of the VAF spectrum, as well as mosaics that were flagged by MosaicHunter quality filters. Candidates mosaic variants were considered confirmed by MiSeq analyses if they demonstrated an amplicon VAF exceeding 0.01 but less than 0.45, so as to indicate a variant of post-zygotic origin. MiSeq VAF values closely correlated with those originally determined by exome sequencing (*P*= 2.2×10^-16^**; Fig S5**).

We confirmed 85/97 (88%) EM-mosaic candidate mosaic variants. Three candidate variants were likely germline *de novo* SNVs (VAF>0.45). Nine candidate variants were ‘false positives’ that were neither germline *de novo* SNVs or mosaic SNVs since either no variant reads were detected by MiSeq sequencing of the proband amplicon, or the same small fraction of variants were detected in proband amplicon and one parent’s amplicon.

Parallel analyses with MosaicHunter confirmed 44/68 (65%) candidate mosaic variants. There were 23 sites for which no variant reads were detected by MiSeq amplicon sequencing (MiSeq VAF<0.001) or in which the same small fraction of variant reads was detected in the proband amplicon as in one parent’s amplicon.

We considered whether estimates of mosaic variant frequency were sensitive to whole exome sequencing depth by calibrating estimates of mosaic detection power using properties of the sequence data (average read depth, prior mosaic fraction, and the value of our overdispersion parameter *θ*) (**Fig S6; Supplemental Methods**). Our projected mosaic detection power curves demonstrated more than a doubling of power to detect mosaic variants with VAF 0.2 as sequencing depth increases from 40x to 80x (**Fig 1C**). Projected mosaic detection power curves for less stringent mosaic cutoffs showed similar increases of power with increasing sequencing depth (**Fig S8**).

To estimate the ‘true’ frequency of mosaicism per blood DNA exome, independent of average coverage detection power constraints, we estimated the ‘true’ mosaic count in a VAF range by multiplying the number of mosaics by the inverse of the detection power for each VAF bin. Applying this method to the 184 of 309 high-confidence EM-mosaic variants with VAF>0.1, we estimated the adjusted number of mosaics with VAF>0.1 to be 361 (**Fig S8A**). Thus, the true frequency of coding mosaics in the blood (0.4>VAF >0.1) is 0.14 variants per individual, representing a non-negligible class of mutations with potential contribution to genetic risk for congenital heart disease. The estimated true mosaic frequency does not change significantly when using less stringent mosaic definitions (**Figure S8**). In sum, we identified 315 blood mosaic variants in 2530 CHD probands or 0.13 mosaic variants per subject with a mean VAF of 0.13±0.06. We do not anticipate that doubling the sequencing depth would change significantly this estimate.

### Mosaic variants occurred most frequently at CpG sequences

Previous studies demonstrated a strong preference for *de novo* C>T mutations at CpG dinucleotides compared to other dinucleotides due to the spontaneous deamination of 5-methylcytosine {Fryxell 2005; Francioli 2015}. We asked whether the germline *de novo* variants observed in CHD probands and the 332 mosaic sites demonstrated a similar sequence preference (**Fig 1, Table 1, S3, S4**). Of the 2662 germline *de novo* mutations identified in 2530 CHD probands, 979 variants (37% of all variants) involved mutation of the cytosine of a CpG dinucleotide (**Fig 2A**). By contrast, 99 (29% of all mosaic SNVs) of 332 mosaic SNVs altered the cytosine of a CpG dinucleotide; significantly more than expected by chance (2.2x above expectation; p=2×10^-15^). These observations suggest that somatic *de novo* mutations were 1.4-fold less likely to involve a CpG dinucleotide than germline *de novo* mutations in CHD probands (*P*=0.01; **Fig 2B**). Even ignoring the high CpG mutation frequency, cytosines and guanines were ~2-fold more likely to be mutated than adenines or thymidines both for germline mutations and for mosaic variants. Surprisingly, somatic mutations of A>C/T>G transversions in ApC dinucleotides were ~2-fold greater than the corresponding germline mutations (*P*=5×10^-8^; **Fig 2B**).

**Fig 2.**
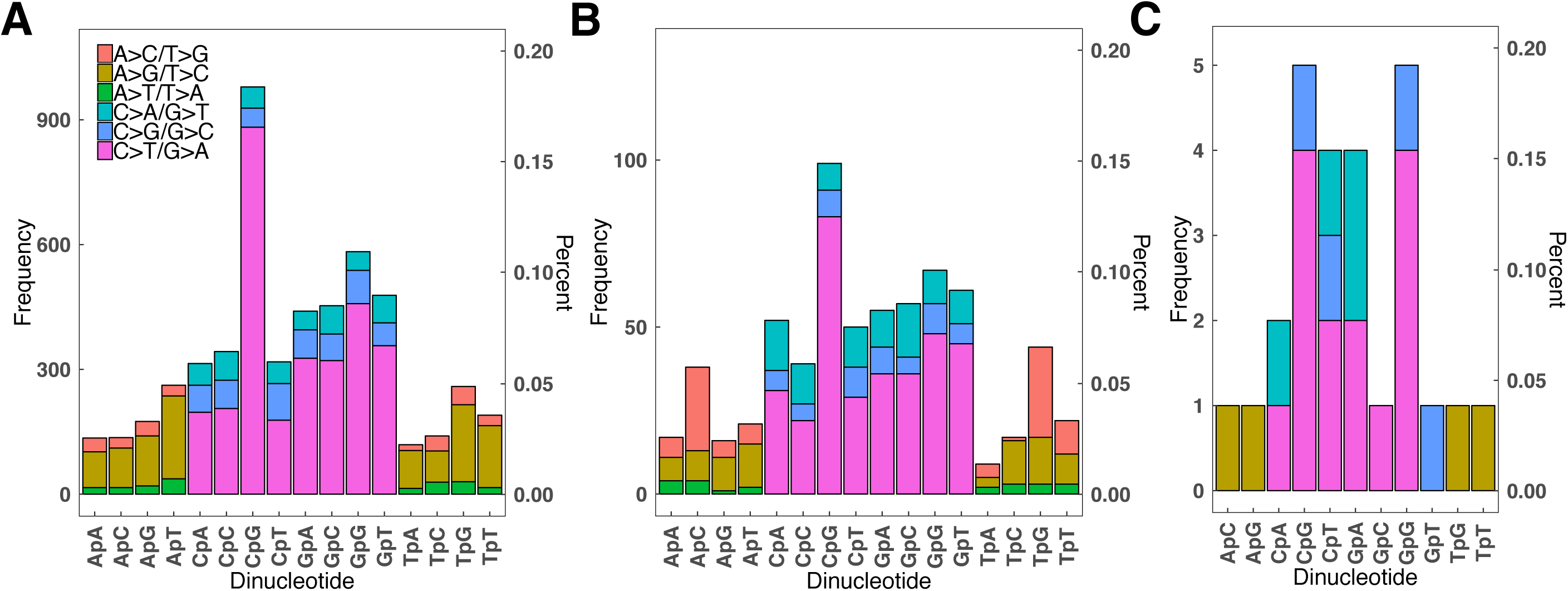
Mutation spectrum of detected germline and mosaic variants. Rates of specific mutations were compared in germline (**A**), blood mosaic (**B**), and CHD tissue mosaic (**C**) variants. Transitions predominated in both variant sets.

**Fig 3.**
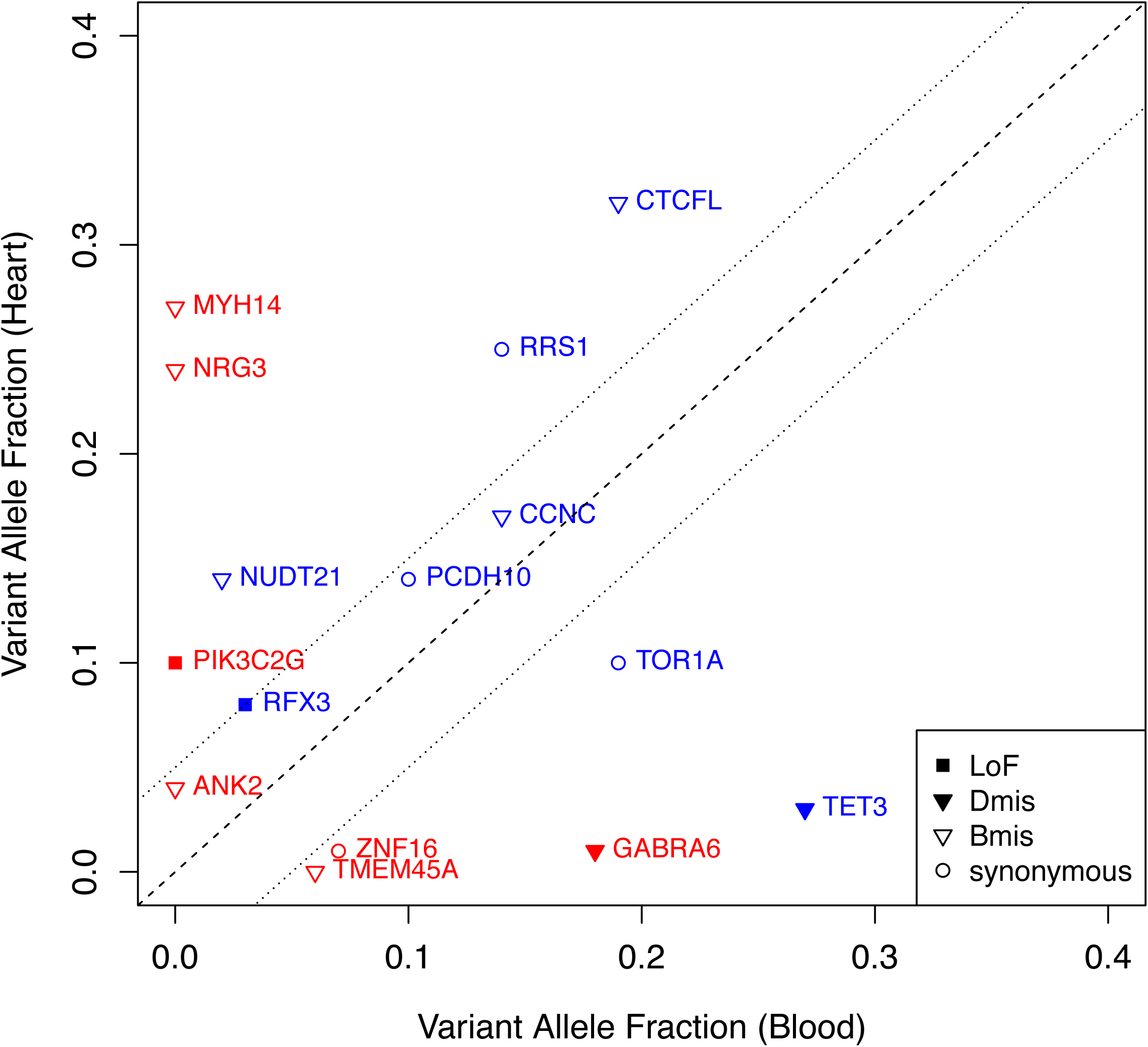
Validated mosaics detected in probands with matched blood and cardiovascular tissue samples available. Validation VAF from blood compared to validation VAF from cardiovascular tissue demonstrated tissue-specific mosaicism (red) as well as shared mosaicism (blue). Predicted effect of mosaic variants corresponds to marker shape.

### Detection of mosaic mutations in CHD tissues

Using EM-mosaic and MosaicHunter we analyzed exome sequences from 70 cardiac tissues derived from 66 subjects with CHD (**Table S6**) and paired blood samples. Among 57 *de novo* variants (allele depth approximately 0.5) that were previously identified in blood-derived DNA, 54 were also found in CHD tissues. Of the 3 *de novo* variants not present in cardiac tissue, 1 was outside of the tissue WES capture region and 2 occurred in a single proband (Table 2). In addition, 23 distinct candidate mosaic variants were detected by EM-mosaic (n=13), MosaicHunter (n=6), or by both algorithms (n= 4). All 23 candidates were tested via MiSeq amplicon sequencing of blood and cardiac tissue DNAs; 15 of 23 unique candidate mosaics were confirmed (**Table 2, S7**), including a *CCNC* variant that was identified in two different CHD tissues from proband 1-01684. Ten (86%) confirmed mosaic variants were detected in blood and cardiac tissues (MAF>0.01), four were found only in cardiac tissue, and one was found only in blood. Of the 7 mosaics detected by blood WES analysis, 4 were confirmed in the corresponding cardiac tissue sample. Remarkably, five confirmed cardiac tissue mosaic variants occurred in one proband (1-07004), one of which was also present in blood DNA.

**Table 2.**
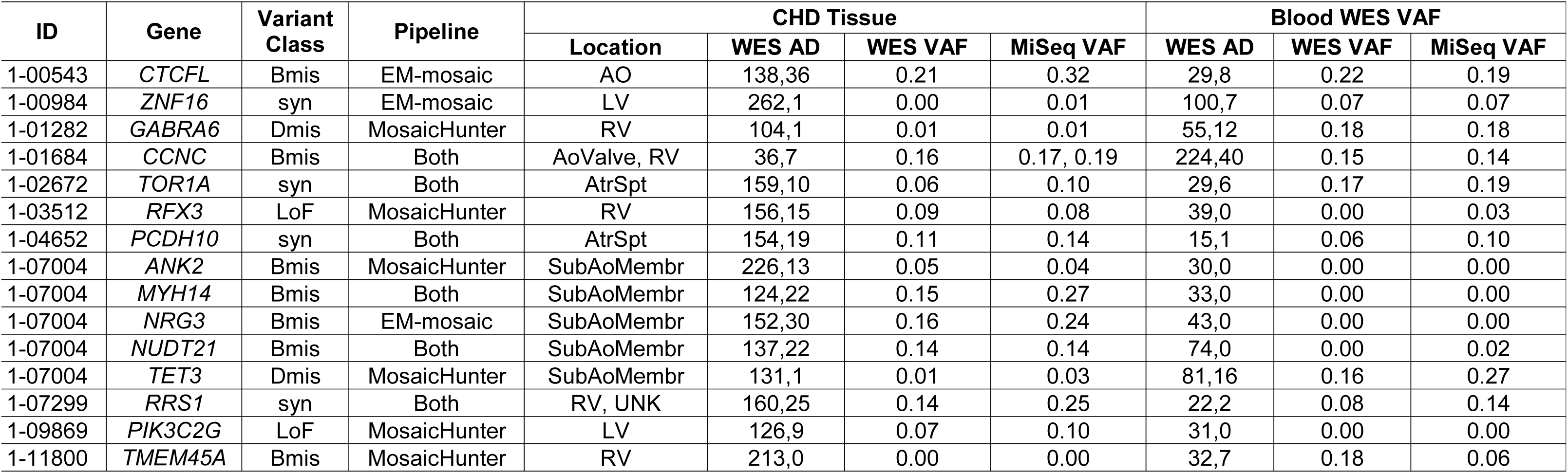
Mosaics detected in individuals with matched cardiovascular tissue and blood. Characteristics of mosaic variants predicted for individuals with blood and cardiovascular tissue WES data available. Among 15 mosaics, 5 were detected via analysis of blood WES, 8 were detected from cardiovascular tissue WES, and 2 were detected by both approaches. Six of 7 (86%) mosaics detected from analysis of blood were present in both DNA sources with MiSeq VAF≥0.01. Two additional variants previously identified as *de novo* germline variants in blood WES were absent from CHD tissue WES. Minimum 1023 MiSeq reads used to determine VAF. Abbreviations: AD, allelic depth (reference, alternate); AO, aorta; AtrSpt, atrial septum; Bmis, benign missense; Dmis, deleterious missense; LOF, Loss of function variant; LV, left ventricle; RV, right ventricle; *VAF, variant allele fraction

These analyses indicate a frequency of coding mosaics (0.4>VAF >0.1) in the cardiac tissues of 0.14 per individual (9 of 66 probands), which approximated our estimate of 0.14 blood mosaics per individual (**Fig S8A**). Despite these similar frequencies, multiple distinct mosaic variants were identified in these tissues. Mosaics with highest VAF were more likely to be found in both tissues (Mann-Whitney U Test *P*=0.019), presumably indicating that the mutation occurred earlier in lineage development (**Fig S9**).

### Blood and Cardiac Tissue Mosaics Likely to Contribute to CHD

Our prior genetic studies of CHD studies showed that damaging *de novo* variants typically occurred in genes highly expressed in the top quartile of the developing E9.5 mouse heart (HHE). {Zaidi 2013; Homsy 2015} or contributed to CHD in mouse models {Jin 2017}. Among the 342 mosaic variants identified from blood or cardiac tissue analyses that were not false by MiSeq, 65 altered these HHE and/or mouse CHD genes (n=4558, **Table S8**). RefSeq functional annotation predicted 52 variants as likely-damaging variants (LOF, Dmis), and 46 as likely benign, missense (**Table S8, S9**). In total, we observed potentially CHD-causing mosaic mutations in 25 participants, representing 1% of the 2530 total participants in our CHD cohort. Among these 25 mosaics, we confirmed 22/22 (100%) candidates tested via MiSeq. Notably, multiple likely-damaging mosaic variants altered genes (*ISL1*, *SETD2*, *NOVA2*, *SMAD9*, *LZTR1*, *KCTD10*, *KCTD20*, *FZD5,* and *QKI)* involved in key developmental pathways, which may account for the extra-cardiac phenotypes observed in these patients (**Table 3, S10**). There was no difference in the proportion of individuals with extracardiac features among those with damaging mosaic variants compared to the overall cohort (11/25 vs 909/ 2521, *P*=0.68), and there was a wide range of CHD subtypes. Five subjects carried additional *de novo* LoF or Dmis variants (1-06216, *TYRP1*; 1-04046, *KRT13*; 1-06677, *TRIP4;* 1-05011, *KDM5B*; 1-00018, *SBF1*) and 4 genes harbored *de novo* LoF or Dmis variants other than those listed in Table 3 (*FBN1*; *PKD1*; *LZTR1*; *PIK3C2G*). No CNVs were detected in these subjects, with the exception of 1-00192 (duplication at chr15:22062306-23062355; non-overlapping with the *GLYR1* mosaic).

**Table 3.** Damaging Mosaics in CHD-relevant genes. There were 25 potentially-pathogenic mosaic mutations based on known gene function and patient phenotype. Some of these probands have previously described rare LoF/Dmis variants, though none are likely pathogenic for CHD {Jin 2017}. Additionally, some genes were previously found to have LoF/Dmis variants among other individuals in this CHD cohort. Abbreviations: ASD, atrial septal defect; BAV, bicuspid aortic valve; Dmis, deleterious missense; episcore, haploinsufficiency score (percentile rank) {Han 2018}; Heart Exp, heart expression percentile rank; LoF, loss-of-function; pLI, probability of loss-of-function intolerance {gnomAD}; PCGC, Pediatric Cardiac Genomics Consortium; VAF, variant allele fraction; VSD, ventricular septal defect. *VAF refers to CHD tissue WES.

| ID | Gene | Variant Class | Blood VAF | pLI | Episcore | Heart Exp | Age (y) | Clinical Phenotype |  | PCGC denovo LoF/Dmis Variants in Mosaic Gene |
| --- | --- | --- | --- | --- | --- | --- | --- | --- | --- | --- |
|  |  |  |  |  |  |  |  | Cardiac Abnormalities | Extracardiac Abnormalities |  |
| 1-00761 | *FBN1* | Dmis | 0.24 | 1.00 | 98 | 93 | 4.3 | Mitral stenosis | dysmorphic features, subglottic stenosis, hypoplastic left mainstem bronchus, short stature | 3 |
| 1-07004 | *TET3* | Dmis | 0.16 | 1.00 | 7 | 87 | 10.3 | Subaortic stenosis | None | 0 |
| 1-05662 | *SETD2* | LoF | 0.13 | 1.00 | 99 | 85 | 0.8 | Aortic coarctation, mitral valve hypoplasia | None | 0 |
| 1-00344 | *UBR5* | splice | 0.27 | 1.00 | 95 | 90 | 16 | D-transposition of the great arteries, VSD, valvar and subvalvar pulmonary stenosis | None | 0 |
| 1-03512 | *RFX3* | LoF | 0.09 | 1.00 | 100 | 46 | 0.4 | Tetralogy of Fallot with pulmonary stenosis | None | 0 |
| 1-06216 | *ITSN1* | Dmis | 0.21 | 1.00 | 98 | 86 | 0.3 | ASD | plagiocephaly, rib anomaly, single kidney, dysmorphic facial features | 0 |
| 1-00363 | *QSER1* | Dmis | 0.06 | 1.00 | 94 | 79 | 3.7 | Tetralogy of Fallot with pulmonary stenosis, VSD | inguinal hernia | 0 |
| 1-13185 | *PKD1* | Dmis | 0.10 | 1.00 | 87 | 84 | 0.8 | VSD, partially anomalous pulmonary venous return | hemangioma | 1 |
| 1-00192 | *GLYR1* | Dmis | 0.22 | 0.99 | 89 | 93 | 0.4 | ASD, VSD, interrupted aortic arch, hypoplastic tricuspid valve, BAV | None | 0 |
| 1-04046 | *FZD5* | Dmis | 0.09 | 0.99 | 89 | 48 | 0.2 | Tetralogy of Fallot with pulmonary stenosis, VSD | None | 0 |
| 1-06649 | *NOVA2* | Dmis | 0.15 | 0.95 | 75 | 56 | 0.6 | Tetralogy of Fallot with pulmonary stenosis | None | 0 |
| 1-05095 | *ISL1* | LoF | 0.07 | 0.90 | 97 | 25 | 2.4 | ASD | None | 0 |
| 1-06677 | *KCTD10* | Dmis | 0.16 | 0.84 | 75 | 91 | 10.1 | Aortic coarctation, pulmonary valve stenosis | dysmorphic facial features, hydrocephalus, pyloric stenosis, single kidney, imperforate/atretic anus | 0 |
| 1-05447 | *HNRNPAB* | Dmis | 0.09 | 0.76 | 72 | 99 | 7.8 | ASD, BAV, aortic coarctation | None | 0 |
| 1-00021 | *QKI* | LoF | 0.13 | 0.76 | 94 | 97 | 0.5 | Doublet outlet right ventricle, pulmonary stenosis, VSD | None | 0 |
| 1-11871 | *FHOD3* | Dmis | 0.18 | 0.05 | 91 | 92 | 0.0 | Tetralogy of Fallot with pulmonary atresia | hypocalcemia, thrombocytopenia, lymphopenia | 0 |
| 1-01458 | *HK2* | Dmis | 0.27 | 0.04 | 89 | 90 | 0.0 | Hypoplastic left heart with aortic and mitral atresia, aortic coarctation | None | 1 |
| 1-00669 | *PRKD3* | splice | 0.19 | 0.02 | 77 | 82 | 0 | D-transposition of the great arteries, conal VSD, bilateral conus, interrupted aortic arch | None | 0 |
| 1-00524 | *RNF20* | LoF | 0.10 | 0.00 | 55 | 83 | 23.7 | Left-dominant complete atrioventricular canal | Heterotaxy with situs inversus totalis, asplenia, duodenal atresia | 0 |
| 1-01851 | *SUCLA2* | LoF | 0.11 | 0.00 | 72 | 89 | 15.5 | Balanced complete atrioventricular canal, aortic coarctation | None | 0 |
| 1-03885 | *LZTR1* | Dmis | 0.20 | 0.00 | 31 | 84 | 14.7 | Abnormal pulmonary vein draining into the right atrium | left sided/midline liver, asplenia, malrotation | 2 |
| 1-05011 | *KCTD20* | Dmis | 0.26 | 0.00 | 76 | 77 | 24.5 | Transposition of the great arteries, Tricuspid and Pulmonary valve atresia | left-sided/midline liver | 1 |
| 1-00018 | *FIG4* | Dmis | 0.19 | 0.00 | 49 | 70 | 11.8 | BAV, mitral atresia, aortic coarctation, VSD, total anomalous pulmonary venous return | nephritis | 1 |
| 1-05661 | *SMAD9* | Dmis | 0.06 | 0.00 | 84 | 39 | 9.3 | Common atrioventricular canal | None | 0 |
| 1-09869 | *PIK3C2G* | LoF | 0.07* | 0.00 | 73 | 28 | 6.3 | Common atrioventricular canal, aortic stenosis, aortic arch hypoplasia, VSDs | dysmorphic facial features, low-set ears, campomelic dysplasia | 1 |

If mosaic variants were unrelated to CHD, we would expect similar allelic fractions between mosaics with variants predicted as likely damaging or likely benign. However, we found that the allele fraction of likely damaging variants was significantly higher (Mann-Whitney U Test *P*=0.001, **Fig 4A**). Moreover, among mosaic variants in genes that are not included among HHE or mouse CHD genes, we found no significant difference of allele fraction (*P*=0.985, **Fig 4B**). We repeated these analyses using less stringent posterior odds cutoffs of 2 and 5 and found the same result (**Fig S10**). Together these data support our conclusion that at least some likely-damaging mosaic variants identified here contribute to CHD. These results were determined independently of MiSeq validation results.

**Fig 4.**
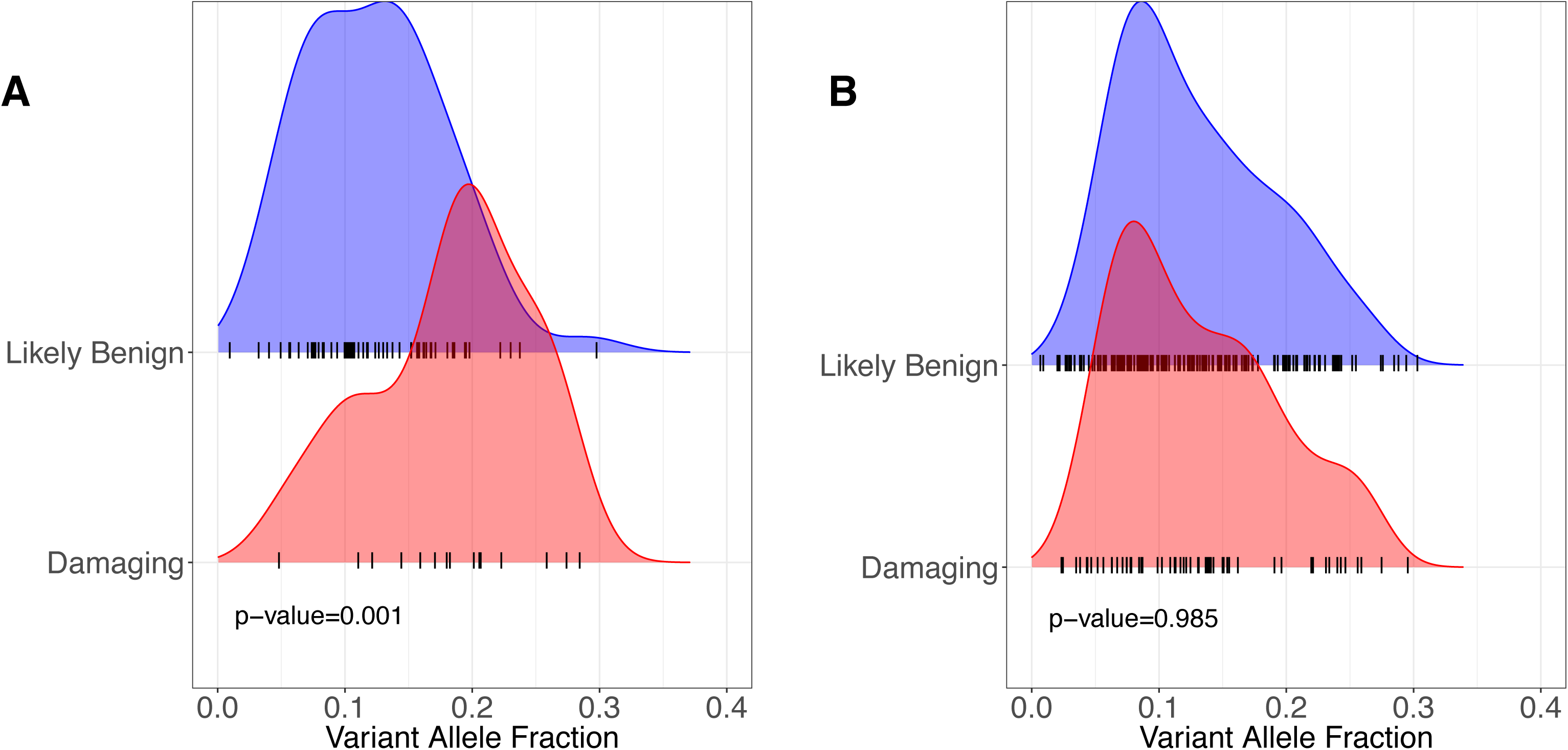
Damaging mosaics in CHD-related genes have higher variant allele fraction than likely-benign mosaics. (**A**) Among the 76 mosaics in CHD-related genes, likely damaging variants have a higher VAF than likely benign (Mann-Whitney *U* p=0.001). (**B**) Among the 233 mosaics in Other (non-CHD-related) genes, there is no difference in VAF based on predicted effect (p=0.985).

## Discussion

Distinguishing mosaic mutations from constitutional mutations has both clinical management and reproductive implications for proband and parents. Individuals with mosaic mutations are generally clinically less severely affected for conditions that affect multiple parts of the body {Happle 1986; Wallis 1990; Cohn 1990; Etheridge 2011; Donkervoort 2015; Weinstein 2016}. Mutations that occur post-zygotically should have no recurrence risk for the parents and could have a recurrence risk of less than 50% for the proband depending on gonadal involvement. This study is among the first investigations of the role of post-zygotic mosaic mutations in CHD. We developed a new computational method to robustly detect mosaic single nucleotide variants from blood WES data at standard read depth. Applying this method to a cohort of 2530 CHD patients, we detected 309 high-confidence mosaics (with a confirmation frequency of 88% in a subset of variants assessed) or 0.12 variants per proband. Sequencing of cardiac tissue to greater depth identified an additional 8 mosaic variants that had not been detected in blood WES, 6 of which are present in cardiac tissue but not blood. We found significantly more variants per proband in cardiac tissue DNA (0.23 variants per proband) than in blood DNA (0.12 variants per proband; p=0.02). While the increased numbers of mosaic variants in cardiac tissue DNA vs blood DNA may reflect technical differences such as sequencing read depth of cardiac tissue DNA vs blood DNA, it is possible that somatic variation occurs more frequently in cardiac tissue of CHD probands than in their blood. Whether or not there are more cardiac tissue mosaic variants in CHD probands than blood DNA variants, we found 10 mosaic variants among 66 CHD proband cardiac tissues with a higher VAF in tissue than in blood (**Table 2**) and 5 variants among these individuals with a higher VAF in blood than in tissue.

In total, we observed potentially CHD-causing mosaic mutations in 25 participants, representing 1% of the 2530 total participants in our CHD cohort. Among these 25 mosaics, we confirmed 22/22 (100%) candidates tested. We found that in CHD-related genes, likely-damaging mosaic mutations have significantly greater alternative allele fraction than likely-benign mosaics, suggesting that some of these variants contribute to CHD. Comparison of blood and cardiovascular tissues demonstrated tissue-specific mosaic variants, though those variants with a higher VAF were more likely to be shared between tissues. Due to limitations of conventional clinical interpretation for both mosaic and constitutional CHD variants (**Supplemental Methods**), we cannot know with complete certainty which among these 25 variants is pathogenic and instead propose that, among our detected mosaics, the 23 detected from blood WES data provide an estimate of the disease-causing mosaics detectable in blood with standard exome-sequencing read depth. Nine of these variants affect genes known to have a role in cardiac development: *ISL1*, *SETD2*, *NOVA2*, *QKI*, *SMAD9*, *LZTR1*, *KCTD10*, *KCTD20*, and *FZD5*.

The mosaic LOF mutation in *ISL1* is likely to be the cause of CHD in participant 1-05095. *ISL1* is a transcription factor essential to normal cardiac development that regulates expression of *NKX*, *GATA*, and *TBX* family genes {Golzio 2012; Colombo 2018} and controls secondary heart field differentiation and atrial septation {Colombo 2018; Briggs 2012}. *ISL1* deficiency has been shown to lead to severe CHD in mice {Cai 2003; Golzio 2012}. Participant 1-05095 has an isolated atrial septal defect consistent with a secondary heart field defect phenotype {Stevens 2010} and has no other previously reported damaging germline variants in CHD-related genes.

Damaging germline *de novo* variants in CHD subjects are enriched in genes related to chromatin modification and RNA processing {Homsy 2015; Jin 2017}. Three genes with damaging mosaic variants discovered here have related functions. *SETD2* is a histone methyltransferase required for embryonic vascular remodeling {Hu 2010}; it is both sensitive to haploinsufficiency and highly expressed in the heart during development. *NOVA2* is a key alternative-splicing regulator involved in angiogenesis that has been shown to disrupt vascular lumen formation when depleted {Giampietro 2015}. *QKI* encodes an RNA-binding protein that regulates splicing, RNA export from the nucleus, protein translation, and RNA stability {Lauriat 2008}. *QKI* is also highly expressed in the heart during development and has been shown to cause CHD and other blood vessel defects in mice when dysregulated {Noveroske 2002}.

Other damaging mosaic variants affect processes known to be relevant to CHD. *SMAD9* is involved in the TGF-beta signaling pathway. TGF-beta signaling plays a critical role in cardiac development and cardiovascular physiology, leading to pulmonary arterial hypertension and cardiac abnormalities in mice when dysregulated {Drake 2015; Soubrier 2013}. *LZTR1* encodes a member of the BTB-Kelch superfamily that is highly expressed in the heart during development and has been associated with Noonan {Yamamoto 2015; Ghedira 2017} and DiGeorge Syndromes {Kurahashi 1995}, both of which are characterized by CHD. *KCTD10* binds to and represses the transcriptional activity of *TBX5* (T-box transcription factor), which plays a dose-dependent role in the formation of cardiac chambers {Tong 2014}. *KCTD10* is highly expressed in the heart during development and has been shown to produce CHD in mice when dysregulated {Ren 2014}. *KCTD20* is a positive regulator of Akt {Nawa 2013} also highly expressed in the heart during development. *FZD5* is haploinsufficient and encodes a transmembrane receptor involved in Wnt, mTOR, and Hippo signaling pathways and has been shown to play a role in cardiac development {Dawson 2013}.

Finally, two mosaic variants found in cardiac tissue, genes encoding *RFX3* and *PIK3C2G,* may be disease-relevant. *PIK3C2G* is a signaling kinase involved in cell proliferation, survival, and migration, as well as oncogenic transformation and protein trafficking {OMIM: 609001; RefSeq}. The effects of *PIK3C2G* haploinsufficiency during cardiac development has not been characterized. *RFX3* is a highly-constrained ciliogenic transcription factor that leads to pronounced laterality defects {Rasmdell 2005} and disruption of *RFX3* leads to congenital heart malformations in mice {Lo 2011 MGI: 5560494}. Notably the RFX3 LoF variant has a 4-fold higher VAF in cardiac tissue than in blood.

Several investigators, who studied cancer and diseases with cutaneous manifestations, proposed that the VAF correlates with time of mutation acquisition and disease burden {Belickova 2016; Sallman 2016; Happle 1986}. In this study, we used VAF as a proxy for cellular percentage and mutational timing, with increasing VAF corresponding to events occurring earlier in development. Thus, we assume that CHD-causal mosaic events identified in blood-derived DNA occurred during or shortly after the gastrulation process (3^rd^ week of development) {Moorman 2003} in the mesodermal progenitor cells that differentiate into both heart precursor cells (cardiogenic mesoderm) and blood precursor cells (hemangioblasts). We found that in CHD-relevant genes, mosaic sites predicted to be damaging tended to have higher VAF than sites predicted to be likely benign, consistent with the hypothesis that these mutations arose early in fetal development and play significant roles in CHD. However, additional functional studies are necessary to fully assess causality. .

Finally, we recognize that while our method is able to detect a large fraction of mosaic variants in blood, our calibrated estimates for the true number of mosaics suggest there are a non-negligible number of additional mutations that were not identified by our method. At our current average sequencing depth of 60x, we have limited sensitivity in the low VAF (<0.05) range. To reliably identify these low allelic fraction sites, ultra-deep sequencing will be critical to distinguishing true variants from noise. At 500x, we estimate detection sensitivity for mosaic events at VAF 0.05 to be above 80%. We also recognize age-related clonal hematopoiesis {Jaiswal 2014; Genovese 2014} as a potential confounding factor in somatic mutation detection; however, our study cohort includes mostly pediatric cases and we did not observe mosaic mutations in genes related to clonal expansion (e.g. *ASXL1, DNMT3A, TET2, JAK2*) nor did we observe a relationship between proband age and mosaic rate (**Fig S11, S12**), suggesting minimal impact from this process.

## Conclusions

This study is among the first investigations of the role of post-zygotic mosaic mutations in CHD. Despite limitations in sequencing depth and sample type, EM-mosaic was able to detect 309 high-confidence mosaics with resequencing confirmation in 88% of cases assessed. Using MosaicHunter, an additional 64 candidate mosaic sites were identified, of which 23/46 (50%) candidates from blood DNA and 4/6 (67%) from CHD tissue DNA validated. In total, we observed potentially CHD-causing mosaic mutations in 25 participants, representing 1% of our CHD cohort, and propose that these 25 cases provide an estimate of the disease-causing mosaics detectable in blood with standard exome-sequencing read depth. Additionally, we found that in CHD-related genes, likely-damaging mosaics have significantly greater alternative allele fraction than likely benign mosaics, suggesting that many of these variants cause CHD and occurred early in development. In the subset of our cohort for which cardiovascular tissue samples were available, we show that mosaics detected in blood can also be found in the disease-relevant tissue and that, while the VAF for mosaic variants often differed between blood and cardiovascular tissue DNA, variants with higher VAF were more likely to be shared between tissues. Given current limitations in sequencing depth and on the availability of relevant tissues, particularly for conditions impacting internal organs like the heart, the full extent of the role of mosaicism in many diseases remains to be explored. However, as datasets containing larger numbers of blood and other tissue samples sequenced at higher depths become increasingly available, we will be able to more fully characterize the biological processes underlying post-zygotic mutation and, by extension, the contribution of mosaicism to disease using the methods presented here.

## Methods

### Samples and Sequencing Data

We analyzed WES data from 2530 Congenital Heart Disease (CHD) proband-parents trio families who were recruited as part of the Pediatric Cardiac Genomics Consortium (PCGC) study {Homsy 2015; Jin 2017}. Genomic DNA from venous blood or saliva was captured using Nimblegen v.2 exome capture reagent (Roche) or Nimblegen SeqCap EZ MedExome Target Enrichment Kit (Roche) followed by Illumina DNA sequencing (paired-end, 2×75bp) {Jin 2017, Zaidi 2013}. Genomic DNA from 70 surgically-discarded cardiovascular tissue samples (2-10mg) was isolated using DNeasy Blood & Tissue Kit (QIAgen), then captured using xGen Exome Research Panel v1.0 reagent (IDT) followed by Illumina DNA sequencing (paired-end, 2×75bp). Sequence reads were mapped to the hg19 human reference genome with BWA-MEM and BAM files were further processed following GATK Best Practices, which included duplication marking, indel realignment, and base quality recalibration steps. Blood and saliva samples had sample average depth 60x and cardiovascular tissue samples had sample average depth 160x.

### De Novo Variant Calling and Annotation

We processed our sample BAMs and called variants on a per-trio basis using SAMtools (v1.3.1-42) and BCFtools (v1.3.1-174). Pileups were generated using samtools ‘*mpileup’* command with mapQ 20 and baseQ 13 to minimize the effect of poorly mapped reads on variant allele fraction, followed by bcftools ‘*call’* using a cutoff of 1.1 for the posterior probability of the homozygous reference genotype parameter (-p) to capture additional sites with variant allele fraction suggestive of post-zygotic origin that would otherwise be excluded under the default threshold of 0.01. To identify *de novo* mutations from trio VCF files, we selected sites with (i) a minimum of 6 reads supporting the alternate allele in the proband and (ii) for both parents, a minimum depth of 10 reads and 0 alternate allele read support. Variants were then annotated using ANNOVAR (v2017-07-17) to include information from refGene, gnomAD (March 2017), 1000 Genomes (August 2015), ExAC, genomicSuperDups, CADD (v1.3) COSMIC (v70), and dbSNP (v147) databases, as well as pathogenicity predictions from a variety of established methods included as part of the dbNSFP (v3.0a) database or generated in-house (MCAP, REVEL, MVP, MPC). We used REVEL {Ionnidis 2016} to evaluate missense variant functional consequence, using the recommended threshold of 0.5 corresponding to sensitivity of 0.754 and specificity of 0.891. We used spliceAI {Jaganathan 2019} to predict the variant functional impact on splicing using the delta score thresholds of 0.2 for likely pathogenic (high recall), 0.5 for pathogenic (recommended), and 0.8 for pathogenic (high precision). We considered sites predicted to be Likely Gene-Disrupting (LOF) (stopgain, stoploss, frameshift indels, splice-site), Deleterious Missense (Dmis; nonsynonymous SNV with REVEL>0.5), or splice-damaging (Benign Missense or synonymous SNV with delta score > 0.5) to be damaging and likely disease causing. We considered sites predicted to be Synonymous (delta score ≤ 0.5) or Benign missense (Bmis; nonsynonymous SNV with REVEL ≤ 0.5 and delta score ≤ 0.5) to be non-damaging.

### Pre-processing and QC

To reduce the number of low VAF technical artifacts introduced by our variant calling approach, we pre-processed our variants using a variety of filters. We first excluded indels from further analysis, as their downstream model parameter estimates were less stable than those of SNVs. We then filtered our variant call set for rare heterozygous coding mutations (Minor Allele Frequency (MAF) ≤ 10^-4^ across all populations represented in gnomAD and ExAC databases). To account for regions in the reference genome that are likely to affect read-depth estimates, we removed variant sites found in regions of non-unique mappability (score<1; 300bp), likely segmental duplication (score>0.95), and known low-complexity {Li 2014}. We then excluded sites located in MUC and HLA genes and imposed a maximum variant read depth threshold of 500. We used SAMtools PV4 to exclude sites with evidence of technical issues using a cutoff of 1e-3 for baseQ Bias and Tail Distance Bias and a cutoff of 1e-6 for mapQ Bias. To account for potential strand bias, we used an in-house script to flag sites that have either (1) 0 alternate allele read support on either the forward or reverse strand or (2) p<1e-3 and (Odds Ratio (OR)<0.33 or OR>3) when applying a two-sided Fisher’s Exact Test to compare proportions of reference and alternate allele read counts on the forward and reverse strands. We also excluded sites with cohort frequency>1%, as well as sites belonging to outlier samples (with abnormally high *de novo* SNV (dnSNV) counts, cutoff = 8) and variant clusters (defined as sites with neighboring SNVs within 10bp). Finally, we applied an FDR-based minimum N_alt_ filtering step (**Fig S7**) to control for false positives caused purely by sequencing errors.

### IGV Visualization of Low Allele Fraction de novo SNVs

To reduce the impact of technical artifacts on model parameter estimation, we manually inspected *de novo* SNVs with VAF<0.3 (n=558) using Integrative Genomics Viewer (v2.3.97) to visualize the local read pileup at each variant across all members of a given trio family. We focused on the allele fraction range 0.0-0.3 since this range is enriched for technical artifacts that could potentially impact downstream parameter estimation. Sites were filtered out if (1) there are inconsistent mismatches in the reads supporting the mosaic allele, (2) the site overlaps or is adjacent to an indel, (3) the site has low MAPQ or is not Primary alignment, (4) there is evidence of technical bias (strand, read position, tail distance), or (5) the site is mainly supported by soft-clipped reads.

### Expectation-Maximization to Estimate Prior Mosaic Fraction and Control FDR

Current estimates for the fraction of *de novo* events occurring post-zygotically are unstable due to differences in study factors such as variant calling methods, average sequencing depth, and paternal ages. In order to use this fraction as a prior probability in our posterior odds and false discovery calculations, we reason that this value must be estimated from the data itself. We used an expectation-maximization algorithm to jointly estimate the fraction of mosaics among apparent *de novo* mutations and to calculate a per-site likelihood ratio score. This initial mosaic fraction estimate gives us a prior probability of mosaicism, independent of sequencing depth or variant caller, and allows us to calculate for each variant the posterior odds that a given site is mosaic rather than germline. To control for false discovery among our predicted mosaic candidates, we chose a posterior odds threshold of 10 to restrict FDR to 9.1%.

### Mosaic Mutation Detection Model

To distinguish variant sites that show evidence of mosaicism from germline heterozygous sites, we modeled the number of reads supporting the variant allele (*N_alt_*) as a function of the total site depth (*N*). In the typical case, *N_alt_* follows a binomial model with parameters *N* = site depth and *p* = mean VAF. However, we observed notable overdispersion in the distribution of variant allele fraction compared to the expectations under this binomial model (**Fig S6**). To account for this overdispersion, we instead modeled *N_alt_* using a beta-binomial distribution {Heinrich 2012; Ramu 2013}. We estimated an overdispersion parameter θ for our model as follows: for site depth values *N* in the range 1 to 500, we (1) bin variants by identifying all sites with depth *N*, (2) calculate a maximum-likelihood estimate *θ θ* value using *N* and all *N_alt_* values for variants in a given bin, and (3) estimate a global *θ* value by taking the average of *θ* values across all bins, weighted by the number of variants in each bin. We then used *θ* in our expectation-maximization approach to jointly estimate prior mosaic fraction and to calculate per-site likelihood ratios.

To calculate the posterior odds that a given variant arose post-zygotically, we first calculated a likelihood ratio (LR) of two models: M_0_: germline heterozygous variant, and M_1_: mosaic variant. Under our null model M_0_, we calculated the probability of observing N_alt_ from a beta-binomial distribution with site depth *N*, observed mean germline VAF *p*, and overdispersion parameter θ. Under our alternate model M_1_, we calculated the probability of observing *N_alt_* from a beta-binomial distribution with site depth *N*, observed site VAF *p*=*N_alt_/N*, and overdispersion parameter *θ*. Finally, for each variant, we calculated LR by using the ratio of probabilities under each model and posterior odds by multiplying LR by our E-M estimated prior mosaic fraction estimate. We defined mosaic sites as those with posterior odds greater than 10 (corresponding to 9.1% FDR). We used posterior odds in this context to be able to control for false discovery, but we output similarly valid p-value and likelihood ratio scores for each *de novo* SNV.

### Mutation Confirmation by MiSeq Amplicon Sequencing

Chromosome coordinates were expanded 500 bp upstream and downstream of the candidate mosaic variants in the UCSC Genome Browser. Primer 3 Plus software was used to design forward and reverse primers to generate 150-300 bp amplimers containing the candidate site. PCR reactions consisting of genomic DNA, primers, and Phusion polymerase were amplified by thermal cycling and purified with AMPure XP beads. The purified PCR product was quantified, and 0.5-1.0 ng of product was used to construct Nextera XT libraries according to the protocol published by Illumina. Libraries were purified using AMPure XP beads, and final libraries were quantified and pooled to undergo sequencing through Illumina MiSeq.

We experimentally tested for the presence our predicted post-zygotic sites in the original blood DNA and cardiovascular tissue DNA samples using Illumina MiSeq Amplicon sequencing. The Amplicon Deep Sequencing workflow, optimized for the detection of somatic mutations in tumor samples, offers ultra-high sequencing depth (>1000x) that gives us the resolution to confirm low VAF variants, accurately estimate site VAF, and to distinguish true variant calls from technical artifacts. Mosaic candidates were considered validated if the variant allele matched the MiSeq call and both the mosaic VAF and MiSeq VAF indicated post-zygotic origin (VAF<0.45).

Mosaic candidates were selected for confirmation on the basis of VAF, plausible involvement in CHD (based on predicted pathogenicity and HHE status), and detection method (**Table S11; Table S12**). We sampled mosaics from both ends of the VAF spectrum to evaluate our ability to distinguish high VAF mosaics (VAF>0.2; n=29) from germline variants and to distinguish low VAF mosaics (VAF<=0.1; n=52) from technical artifacts. Confirmation rate across different VAF bins is shown in **Figure S13**. We also selected for confirmation mosaics detected uniquely by either EM-mosaic or MosaicHunter, for the sake of method comparison (**Table 1**).

To examine a potential source of bias in our candidate selection process, we compared the posterior odds distribution of selected candidate mosaics (n=97) against those not chosen (n=212). We found that our tested candidates had lower posterior odds than untested mosaics (mean_tested_=5.382, mean_untested_=7.050, log_10_-scale; Mann Whitney *U P*=0.002) (**Fig 14**), suggesting that our validation rate is not buoyed by testing variants with the strongest evidence of mosaicism. For method development purposes, we intentionally focused on mosaics with lower posterior odds as these fall in the VAF range for which it is most difficult to distinguish germline from mosaic.

### Investigating the relationship between VAF and pathogenicity

We hypothesized that mosaic contribution to disease is positively correlated with cellular percentage and by extension mutational timing. Here, we used variant allele fraction as a proxy for cellular percentage. We grouped mosaics into likely-damaging and likely-benign and compared the distribution of allele fraction in CHD-related genes. We defined likely-damaging variants as: (a) likely gene-disrupting (LOF) variants (including premature stop-gain, frameshifting, and variants located in canonical splice sites), (b) missense variants predicted to be damaging by REVEL {Ioannidis 2016} (with score ≥ 0.5) or (c) missense variants and synonymous predicted to be splice-damaging by spliceAI (with score > 0.5). One of the main findings from previous CHD studies is that damaging *de novo* variants in genes highly expressed in the developing heart (“HHE”, ranked in the top 25% by cardiac expression data in mouse at E14.5 {Zaidi 2013; Homsy 2015}) contribute to non-isolated CHD cases that have additional congenital anomalies or neurodevelopmental disorders. Therefore, we considered the union of HHE genes and known candidate CHD genes {Jin 2017} as CHD-related genes (n=4558). For mosaics in CHD-related genes and for mosaics in other genes, we used a Mann-Whitney *U* Test to compare the VAF distributions of likely-damaging and likely-benign groups.

### Estimated contribution of mosaicism to CHD

We identified likely disease-causing mosaic mutations on the basis of predicted pathogenicity and presence in genes involved in biological processes relevant to CHD or developmental disorders. Each mosaic mutation was annotated with gene-specific information, including heart expression percentile, probability of loss-of-function intolerance (pLI) score {Lek 2016}, whether dysregulation causes CHD in mice {Smith 2018; Finger 2017}, and gene function {NCBI RefSeq}. We focused on HHE genes, genes with high pLI (pLI>0.9), genes that cause CHD phenotypes in mice, and genes involved in key developmental processes such as Wnt, mTOR, and TGF-beta signaling pathways. Then, for each patient, we used the clinical phenotype to further prioritize mosaic mutations most likely contributing to that individual’s clinical features. Detailed mutation annotation and clinical phenotypes for the mosaic carriers described above can be found in **Table S10**. We estimate the contribution of mosaicism to CHD as the percentage of individuals carrying likely disease-causing mosaic mutations among all individuals in our CHD cohort.

## Abbreviations

ASD: Autism Spectrum Disorder
CHD: Congenital Heart Disease
dnSNV: *de novo* SNV
Dmis: Deleterious Missense mutation
DP_site_: Total Read Depth at a variant site
DP_sample_: Sample-wide Average Read Depth
ExAC: Exome Aggregation Consortium
FDR: False Discovery Rate
gnomAD: Genome Aggregation Database
HHE: High Heart Expression
LOF: Loss-of-Function
LR: Likelihood Ratio
MAF: Minor Allele Fraction
N: Total Read Depth
N_alt_: Alternate Allele Read Depth
OR: Odds Ratio
PCGC: Pediatric Cardiac Genomics Consortium
pLI: Probability of Loss-of-Function Intolerance
PV4: P-value for strand bias, baseQ bias, mapQ bias and tail distance bias (SAMtools)
SNV: Single Nucleotide Variant
VAF: Variant Allele Fraction
WES: Whole Exome Sequencing

## Supplement

### Supplemental Methods

#### Union with Validated de novo SNVs from Jin et al. Nature Genetics 2017

As part of the PCGC program, Jin et al. previously sequenced and processed a cohort of 2871 CHD probands – including 2530 parent-offspring trios used in this study – to investigate the contribution of rare inherited and *de novo* variants to CHD. They called a total of 2992 proband *de novo* variants, including 2872 SNVs and 118 indels, and Sanger confirmed a subset of the most likely-disease causing variants. Since we processed the same proband-parent trios using different variant calling pipelines, we combined the results of our two approaches to provide a more complete input *de novo* call set for mosaic variant detection.

We first processed our SAMtools *de novo* calls using our upstream filters (*n*=2396 sites passing all filters). We then applied the same upstream filters to the published dnSNVs from Jin et al. (*n*=2650 sites passing all filters) before finally taking the union of these two call sets (*n*=3192). There were 1814 sites in the intersection, with 836 sites unique to the Jin et al. calls and 542 sites unique to our SAMtools calls. After preprocessing, outlier removal, and FDR-based minimum N_alt_ filtering, the remaining 2971 dnSNVs were used as input to our mosaic detection model.

#### Mutation Spectrum Analysis

We compared the mutation spectrum – the frequencies of all possible base changes – of our predicted mosaic candidates against the spectrum of our predicted germline heterozygous variants. Under the assumption that that post-zygotic events occur randomly (i.e. due to errors in DNA replication rather than a specific biological process), the mosaic mutation spectrum should not differ significantly from the germline mutation spectrum. We used Pearson’s Chi-square Test to test for a difference in frequencies across all base changes between our predicted sets of variants. We interpreted large qualitative differences in base change frequencies as evidence of technical artifacts and rejection of the Chi-square null as evidence of systemic issues in our pipeline.

#### Mosaic Detection Power Given Sample Average Coverage

To model statistical power in the context of mosaic variant detection, we considered two conditional probabilities: (i) the probability of detecting a mosaic event (i.e. the probability of a variant’s posterior odds exceeding a threshold) given site depth *DP_site_*, VAF, and overdispersion parameter *θ* and (ii) the probability of observing site depth *DP_site_*, given sample-wide average coverage *DP_sample_*.

i. Pr(detect mosaic | *DP_site_*, VAF, *θ*) was calculated by first identifying the VAF range (and by extension, the range of *N_alt_*) over which posterior odds > cutoff, then by integrating the beta-binomial probability mass function over this range, with considerations for the probability of strand bias (P(strand bias | *DP_site_*) ~ Binomial(*N_alt_*, *DP_site_*, *p*=0.5)).
ii. Pr(*DP_site_* | *DP_sample_*) follows an overdispersed poisson distribution that we approximated using a negative binomial model with overdispersion parameter *θ* {Sampson 2011}. For each *DP_sample_* value, we calculated a vector of weights corresponding to Pr(*DP_site_* | *DP_sample_*) for *DP_site_* values in the range (1, 1500).

Finally, we took the sum of the detection probabilities described in (i) multiplied by the weights described in (ii) to determine the probability of detecting a mosaic variant given a sample average coverage value – Pr(detect mosaic | *DP_sample_*). Our estimated detection power curves for a range of sample average coverage values typical of exome-sequencing studies are shown in (**Figure 4A**). Our CHD cohort was sequenced to sample average depth of 60x, with prior mosaic fraction=0.121 and estimated *θ* =116.

To estimate the true rate of mosaicism per exome given sample average coverage, we first split our set of predicted mosaics into VAF bins of size 0.05. For each bin above VAF 0.1, we multiplied the number of mosaics by the inverse of the detection power for that given VAF bin to estimate the true count of mosaic variants in that VAF range, assuming full detection power. Since EM-mosaic is underpowered to detect mosaics with VAF < 0.1 in the blood and since this range is enriched for technical artifacts that potentially affect our counts, we did not apply this scaling procedure to these bins to avoid over-inflating our adjusted mosaic rate estimate (**Figure 4B**).

#### Filtering of MosaicHunter Candidate Variants (Fig S3)

MosaicHunter was used to identify candidate mosaic variants from blood exome-sequencing trio data using default settings {Huang 2014}. Filtering of original MosaicHunter candidate variants excluded, in order, any variant present in ExAC (46634), G to T mutations with fewer than N_alt_<10 oxidative indicating DNA damage {Costello 2013} (3995), non-uniquely called sites (4719), germline SNVs previously called by GATK HaplotypeCaller (591), probands with >20 mosaic variants (1490 in 10 probands), mosaic log posterior likelihood ratio <10 (940), variants with >2 parental alternative allele reads (244), variants with gnomAD population frequency > 1e-4 or located in MUC or HLA genes (40).

#### Filtering of cardiovascular tissue Candidate Variants

We used the MosaicHunter pipeline in trio mode to identify candidate variants in WES data from 70 cardiovascular tissue samples (belonging to 66 unique probands). From the list of variants initially reported by the pipeline using default settings, we applied the same filtration steps listed for MosaicHunter candidate variants in blood samples with the exception of the removal of G to T mutations with fewer than 10 alternative allele reads and the mosaic log posterior likelihood ratio <10. Finally, we removed variants that were identified in either parent or had a total read depth <10 in either parent.

#### Clinical interpretation of mosaic variants – limitations

We note that conventional clinical interpretation of mosaic mutations is challenging for several reasons: (i) it is unclear in which tissues each mosaic mutation is expressed (ii) several study participants were very young at time of clinical assessment and many classical disease features may not yet have developed or been noted, and (iii) the absence of additional clinical features does not necessarily rule out a mosaic mutation as being for the cause of the CHD. For the purposes of this study, we selected these mosaic mutations on the basis of predicted pathogenicity and detection in genes involved in biological processes relevant to CHD or developmental disorders

**Fig S1.**
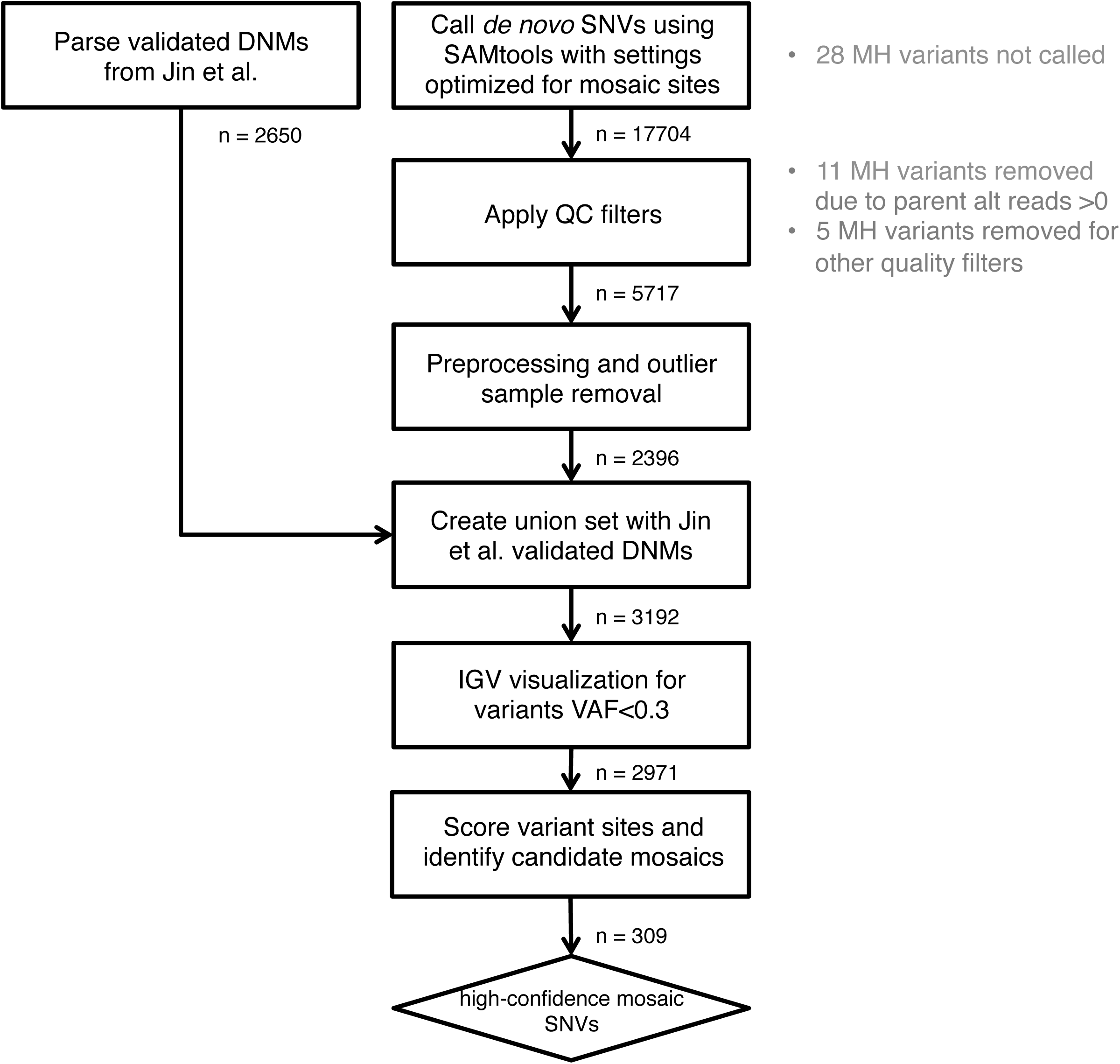
EM-mosaic Flowchart. We first processed our SAMtools *de novo* calls using our upstream filters (*n*=2396 sites passing all filters). We then applied the same upstream filters to the published dnSNVs from Jin et al. (*n*=2650 sites passing all filters) before finally taking the union of these two call sets (*n*=3192). High-confidence mosaics (n=309) were defined as mosaics passing IGV inspection and having posterior odds > 10. Grey text indicates which filters removed candidate mosaic variants called by MosaicHunter but not by EM-mosaic.

**Fig S2.**
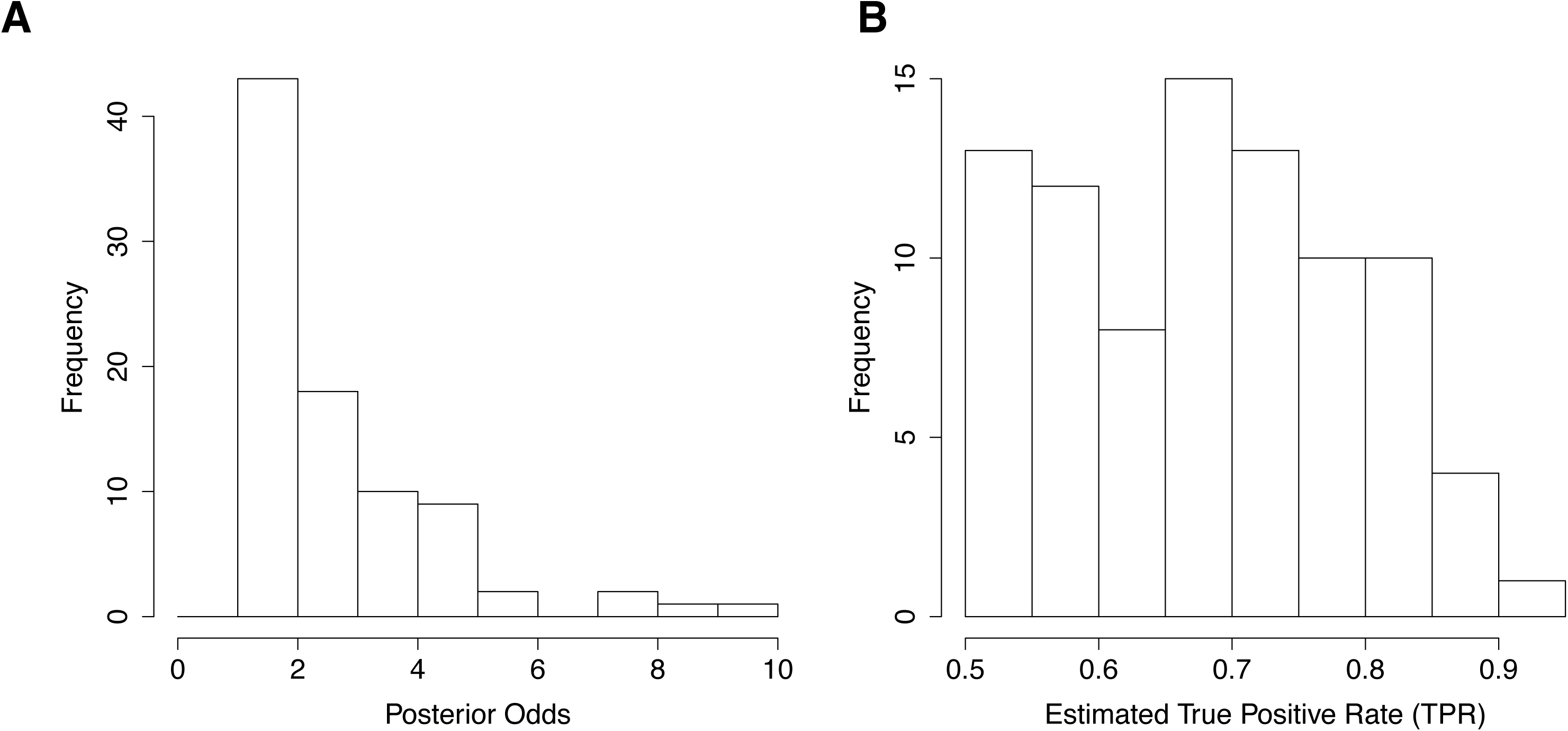
Blood variants with posterior odds between 1 and 10. (**A**) Distribution of the 86 variants with posterior odds between 1 and 10.(**B**) Histogram of counts by bin. To estimate the number of potential mosaics mosaics missed by our threshold, counts of each bin were scaled by the estimated true positive rate (TPR; posterior odds / 1+posterior odds). By our estimate, 54/86 variants were likely mosaic and 32/86 were likely germline.

**Fig S3.**
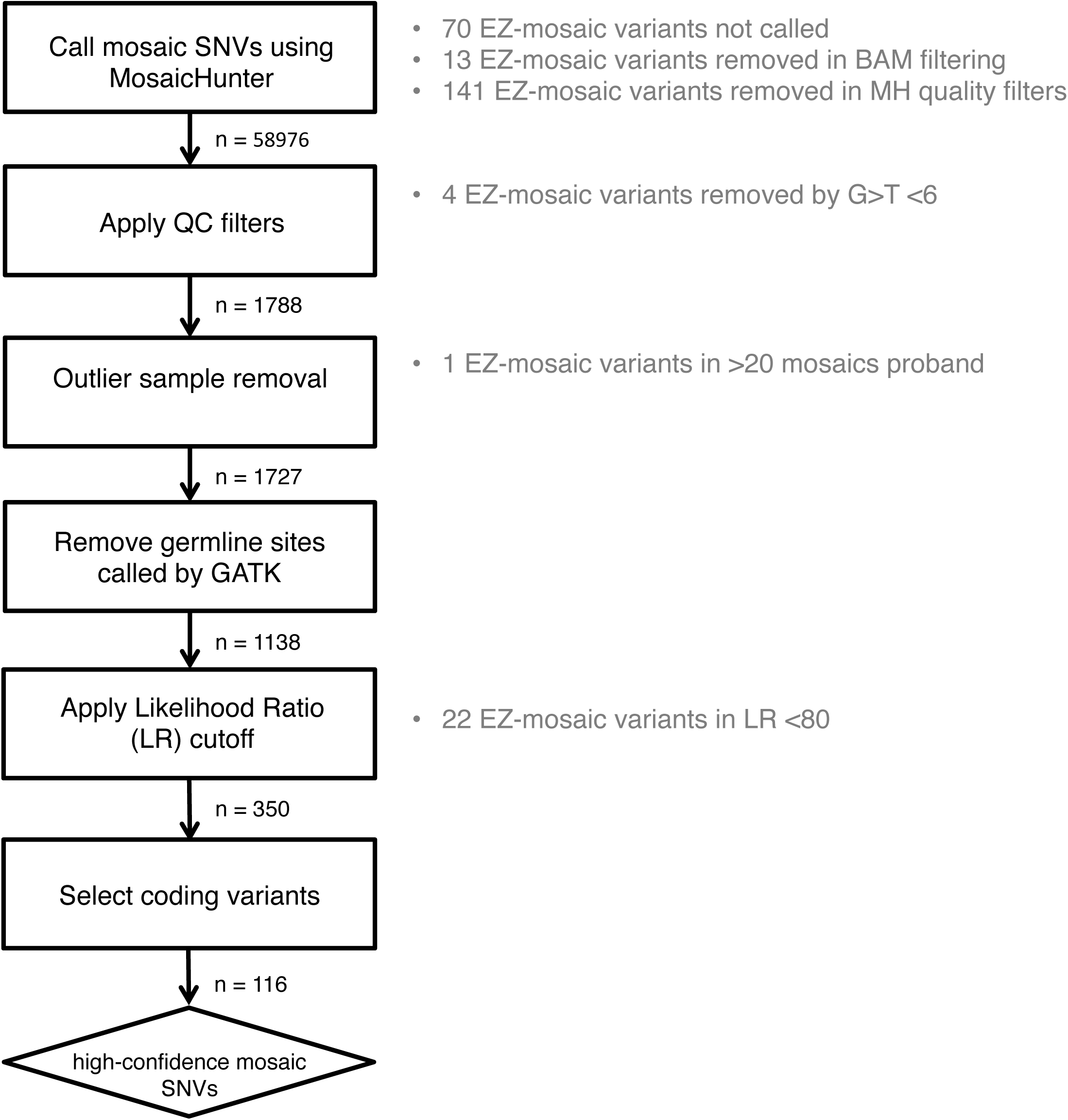
MosaicHunter workflow. Quality Control filters excluded any sites that were (1) present in ExAC (2) G>T with N_alt_<10 (3) parent N_alt_>2. Outliers were defined as probands carrying more than 20 mosaics, or non-unique sites. We also removed sites called as germline by GATK Haplotype Caller. High-confidence mosaics (n=116) were defined as having Likelihood Ratio > 80 and affecting coding regions excluding MUC/HLA genes. Grey text indicates which filters removed variants called by EM-mosaic but not by MosaicHunter.

**Fig S4.**
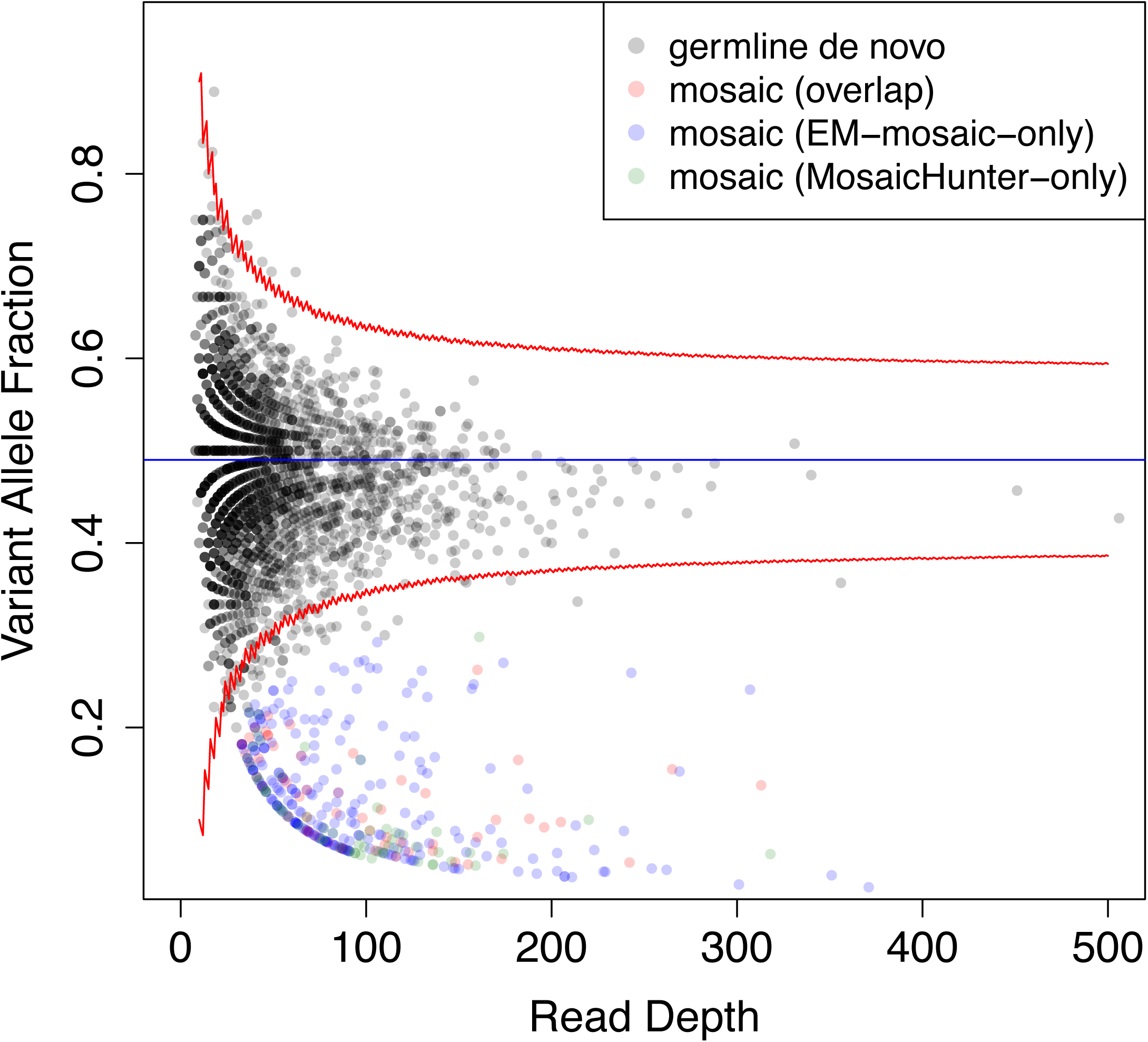
Comparison of variant allele fraction (VAF) and read depth of EM-mosaic and MosaicHunter. Candidate mosiac variants detected by the two pipelines had more overlapping variants at low read depth and VAF values.

**Fig S5.**
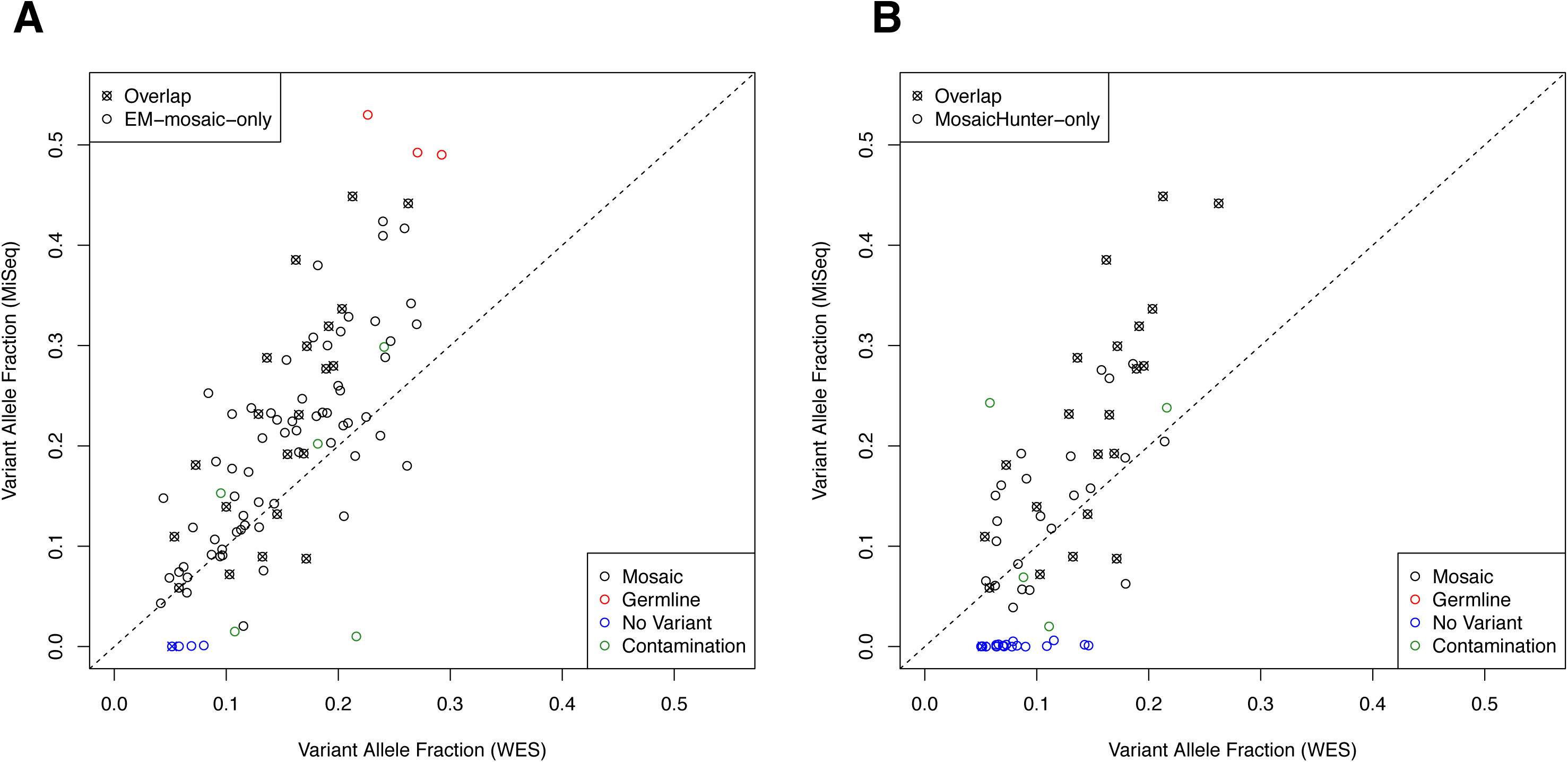
Targeted sequencing to validate candidate blood mosaic variants. (A) EM-mosaic and (B) MosaicHunter variants were assayed using PCR followed by MiSeq for high-depth assessment of mosaicism. Variants with x symbols were shared by both pipelines. Mosaic variants that validated are black, while variants with VAF > 0.45 and therefore germline are red. Validation VAF values demonstrated significant correlation with the original WES-derived VAF for EM-mosaic (Pearson’s correlation *P*=2.2×10^-16^) and MosaicHunter (*P*=8.2×10^-11^).

**Fig S6.**
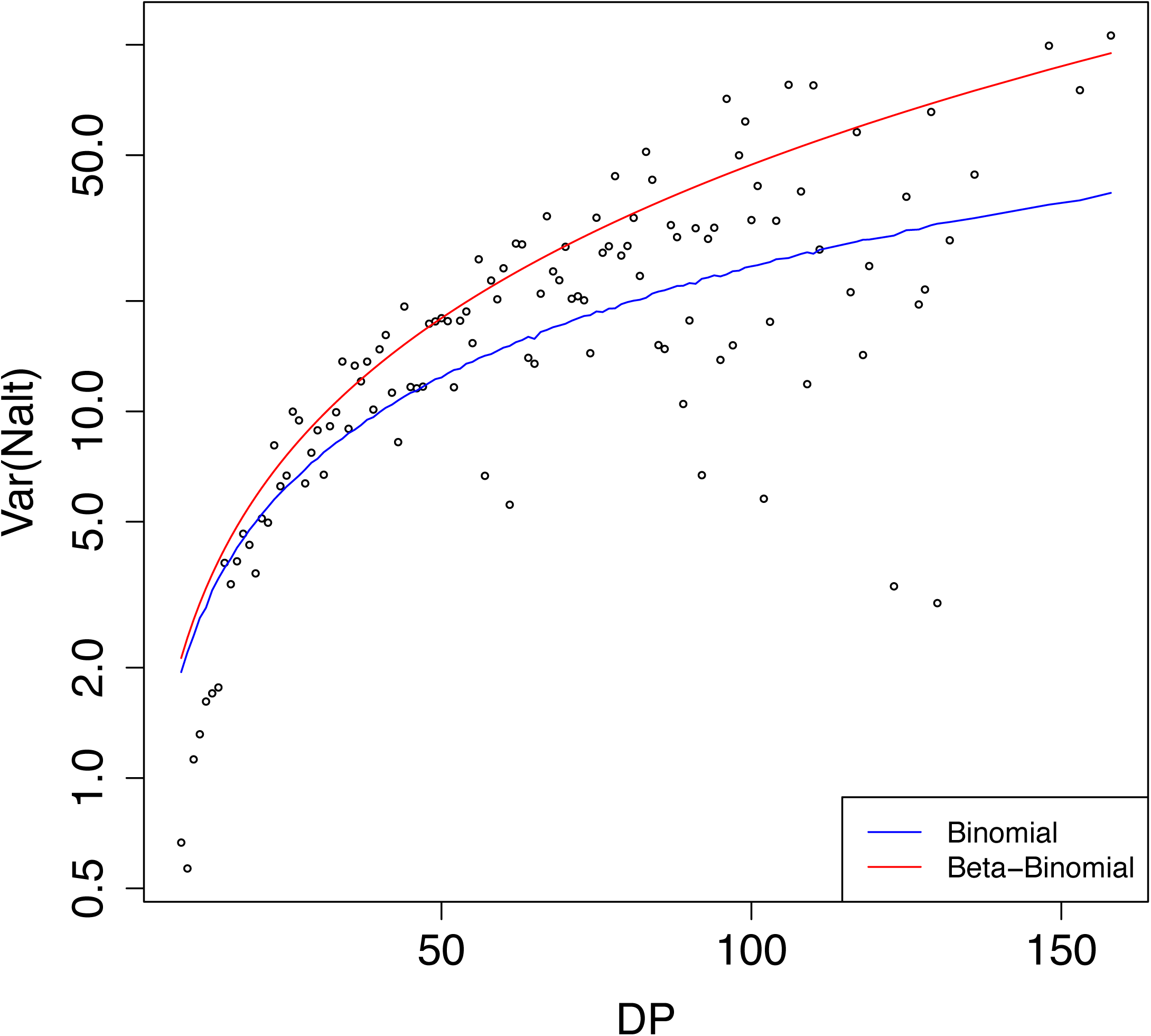
Overdispersion. Overdispersion is commonly seen in WES data {Heinrich 2012; Ramu 2013} and is defined as observing variance (in terms of the VAF of variants with a given DP value) higher than expected across DP values, under a given statistical model. The blue line denotes the expectation under a Binomial model and the red line denotes the expectation under a Beta-Binomial model.

**Fig S7.**
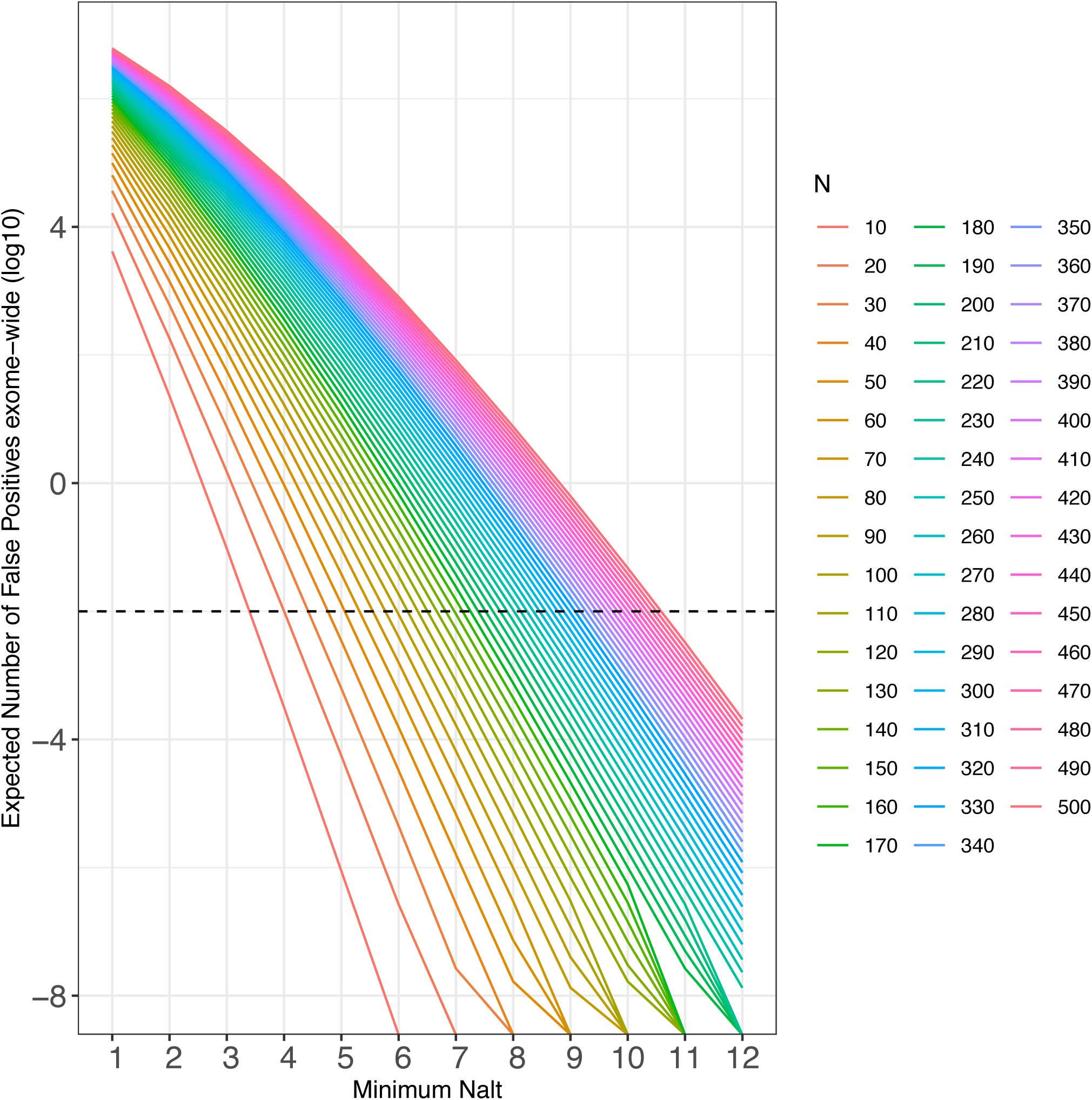
FDR-based minimum N_alt_ threshold. A FDR-based approach was used to determine a threshold for the minimum number of reads supporting the alternate allele for each site to avoid false positives caused purely by sequencing errors. Assuming that sequencing errors are independent and that errors occur with probability 0.005, with the probability of an allele-specific error being 0.005/3=0.00167, and given the total number of reads (*N*) supporting a variant site, we iterated over a range of possible *N_alt_* values between 1 and 0.5*N and estimated the expected number of false-positives due to sequencing error, exome-wide ((1-*f_poisson_*(x=*n*, λ =*N*\*0.005/3))*3×10^7^ ; where *f_poisson_* is the probability of x events in a Poisson process with mean λ). Assuming one coding *de novo* SNV per exome {Acuna-Hidalgo 2016} and that roughly 10% of *de novo* SNVs arise post-zygotically {Lim 2017; Krupp 2017; Freed 2016}, we used a conservative assumption of 0.1 mosaic mutation per exome. To constrain theoretical FDR to 10% we allowed a maximum of 0.01 false positives per exome and used the corresponding *N_alt_* value to define an FDR-based minimum *N_alt_* threshold for each variant. We then excluded variants with alternate allele read counts below this threshold.

**Fig S8.**
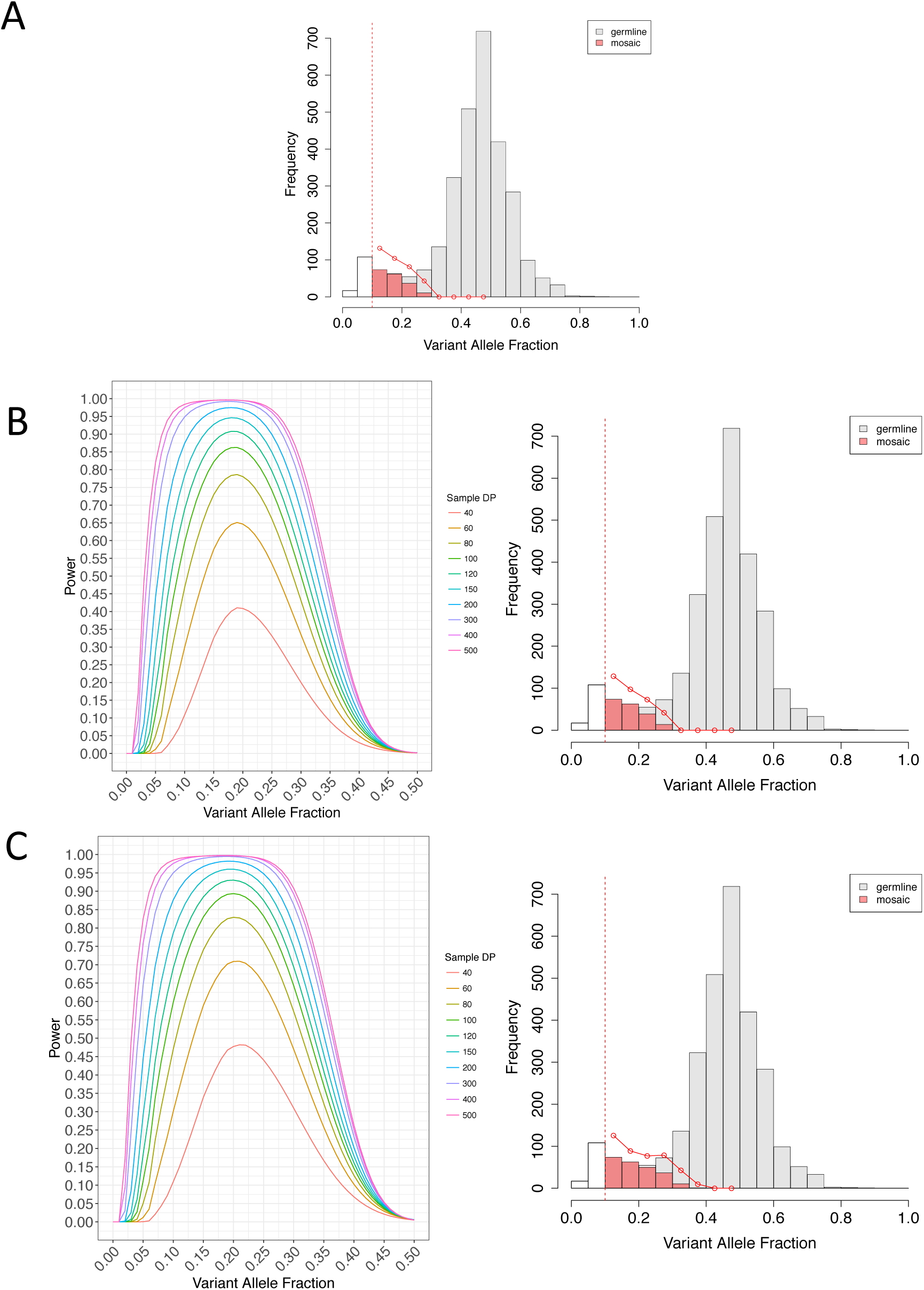
Estimated mosaic detection power using less stringent mosaic definitions. (**A**) Estimated true frequency of detectable coding mosaics (0.4>VAF>0.1) adjusted by detection power (n=341; 0.135/exome) (**B**) Calibrated mosaic detection power and estimated true mosaic frequency of detectable coding mosaics, using posterior odds cutoff of 5 (n=361; 0.143/exome). (**C**) Calibrated mosaic detection power and estimated true mosaic frequency of detectable coding mosaics, using posterior odds cutoff of 2 (0.4>VAF>0.1; n=424; 0.168/exome).

**Fig. S9.**
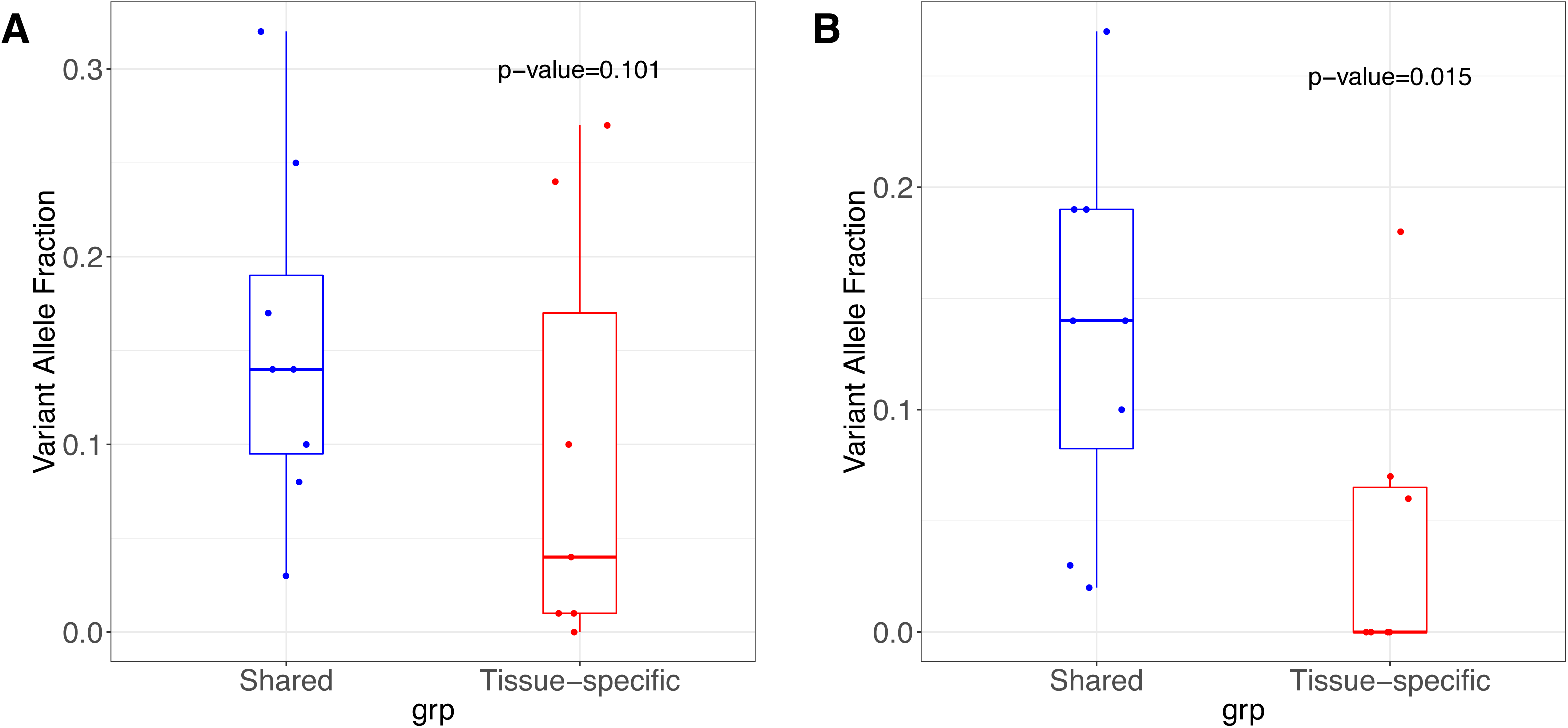
Mosaic variants shared in blood and cardiovascular tissues have higher variant allele fraction. Validation VAF from (**A**) cardiovascular tissue and (**B**) blood had higher VAF for shared variants compared to tissue-specific variants (p=0.101 and 0.015, respectively).

**Fig. S10.**
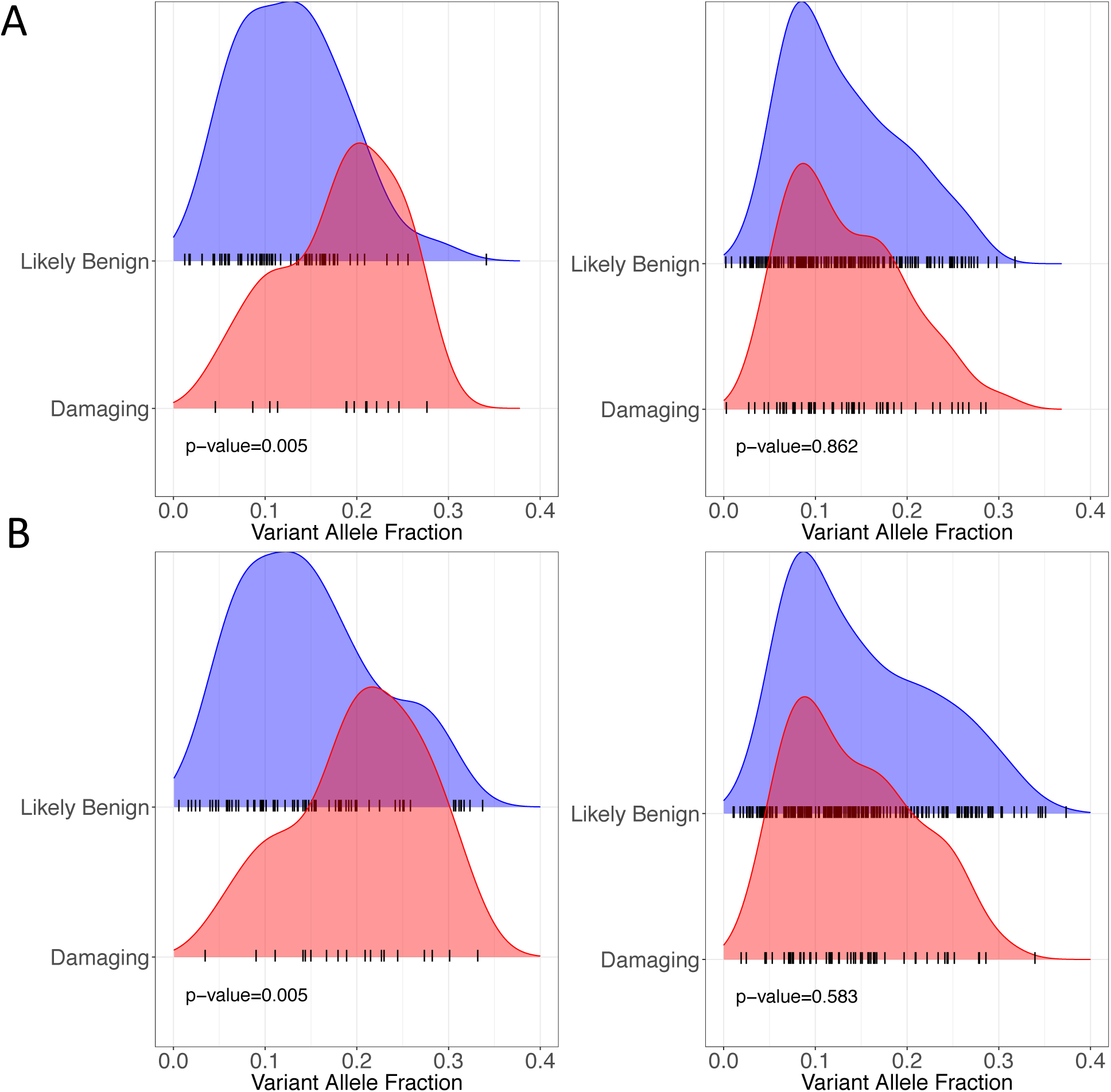
Damaging CHD-related mosaics have higher VAF under less stringent definitions of mosaicism. (**A**) Using posterior odds cutoff of 5 (corresponding to 315 mosaics). Among 78 mosaics in CHD-related genes (left), there were 14 variants predicted as damaging, 63 variants predicted as likely-benign, and 1 variant of unknown functional consequence. Among 237 mosaics in non-CHD-related genes (right), there were 41 variants predicted as damaging, 184 variants predicted as likely-benign, and 2 variants of unknown functional consequence. (**B**) Using posterior odds cutoff of 2 (corresponding to 352 mosaics). Among 89 mosaics in CHD-related genes (left), there were 17 variants predicted as damaging, 71 variants predicted as likely-benign, and 1 variant of unknown functional consequence. Among 263 mosaics in non-CHD-related genes (right), there were 54 variants predicted as damaging, 206 variants predicted as likely-benign, and 3 variants of unknown functional consequence.

**Fig. S11.**
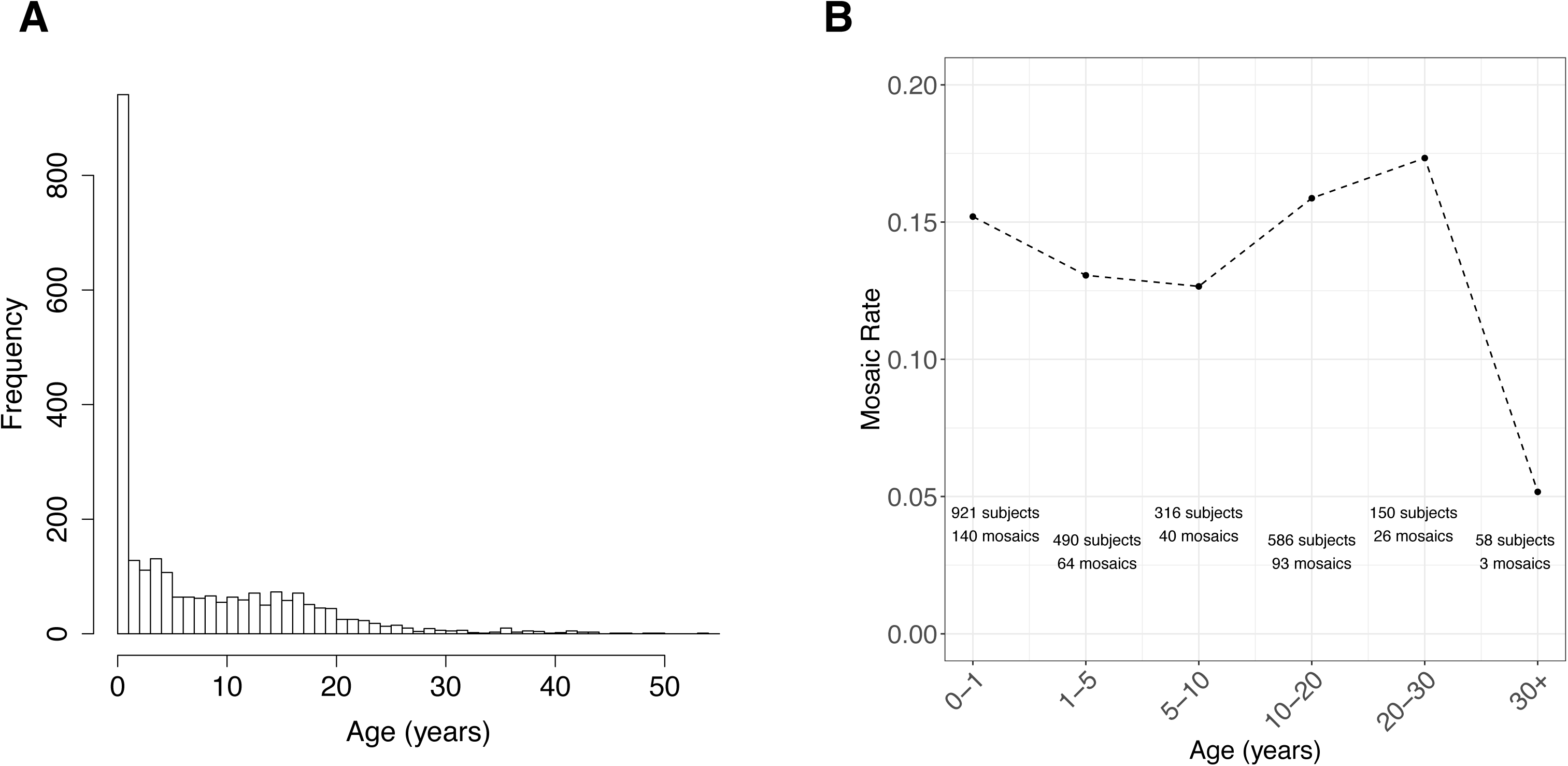
Mosaic rate by age. (**A**) Age distribution for all 2530 probands in cohort. (**B**) Mosaic Rate across Age ranges. Rate = # mosaics/# probands in age bin. Note: 9/2530 probands missing Age information. 1/367 mosaic belong to a proband with missing Age.

**Fig. S12.**
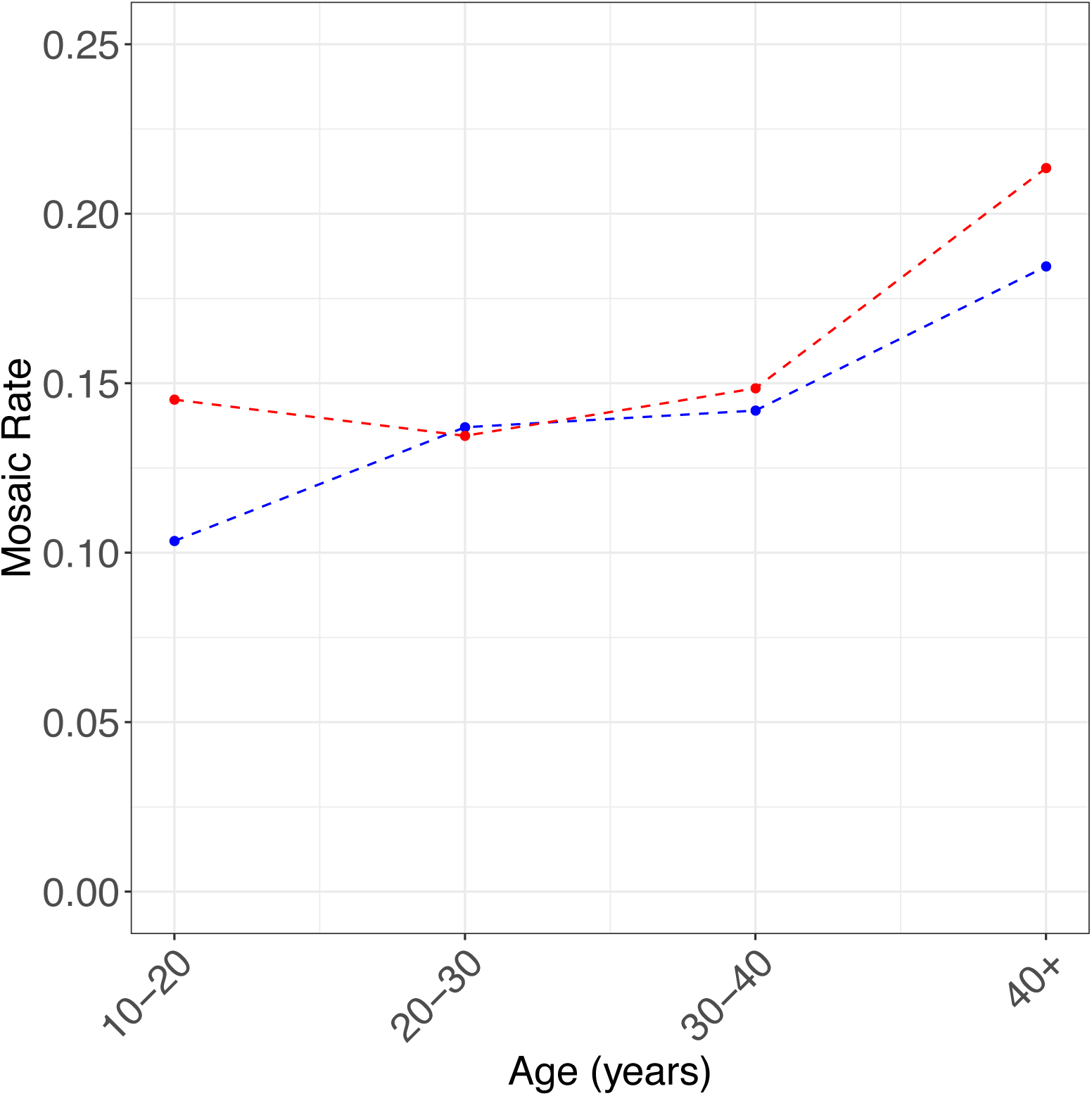
Mosaic rate by parental age at birth. Mosaic rate by age of father (blue) and mother (red) at birth. Rate = # mosaics/# probands in each parental age bin. Note: 9/2530 probands missing age information. 1/367 mosaic belong to a proband with missing age.

**Fig. S13.**
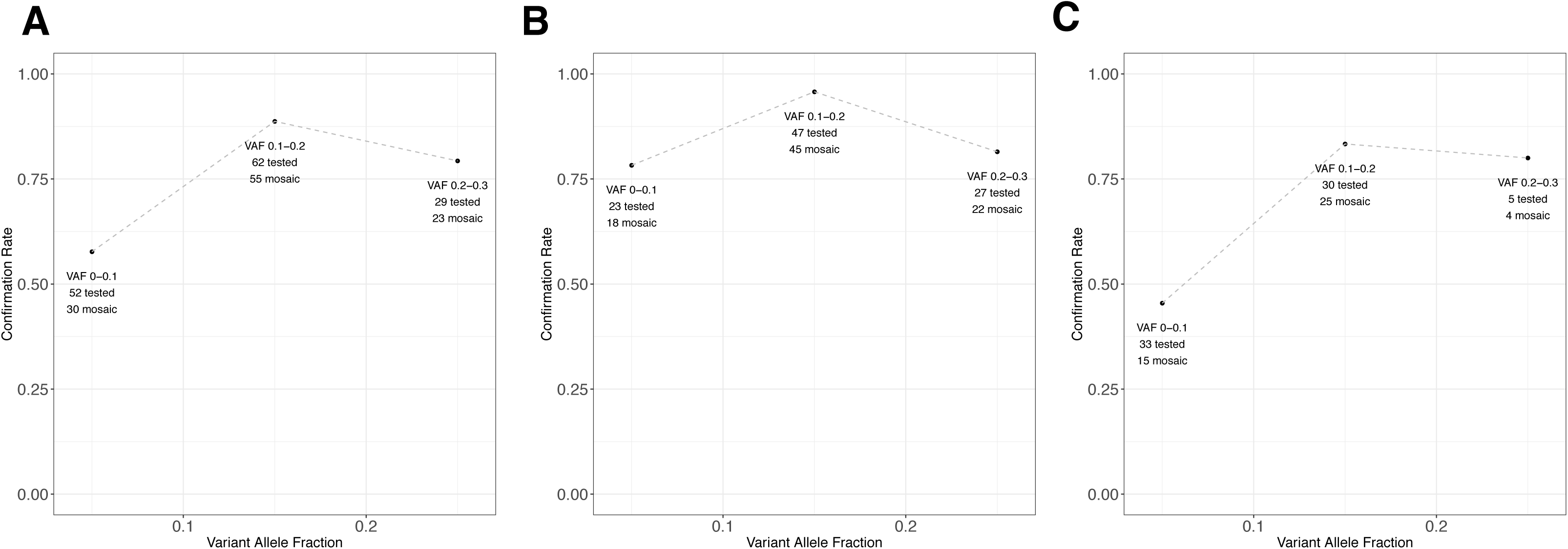
Confirmation rate across VAF bins. The number of candidates for which we performed MiSeq resequencing among (**A**) the union set (n=143 tested) (**B**) all EM-mosaic calls (n=97) and (**C**) all MosaicHunter (n=68) calls vs. the number confirmed as mosaic for VAF ranges [0, 0.1), [0.1, 0.2), and [0.2, 0.3).

**Fig. S14.**
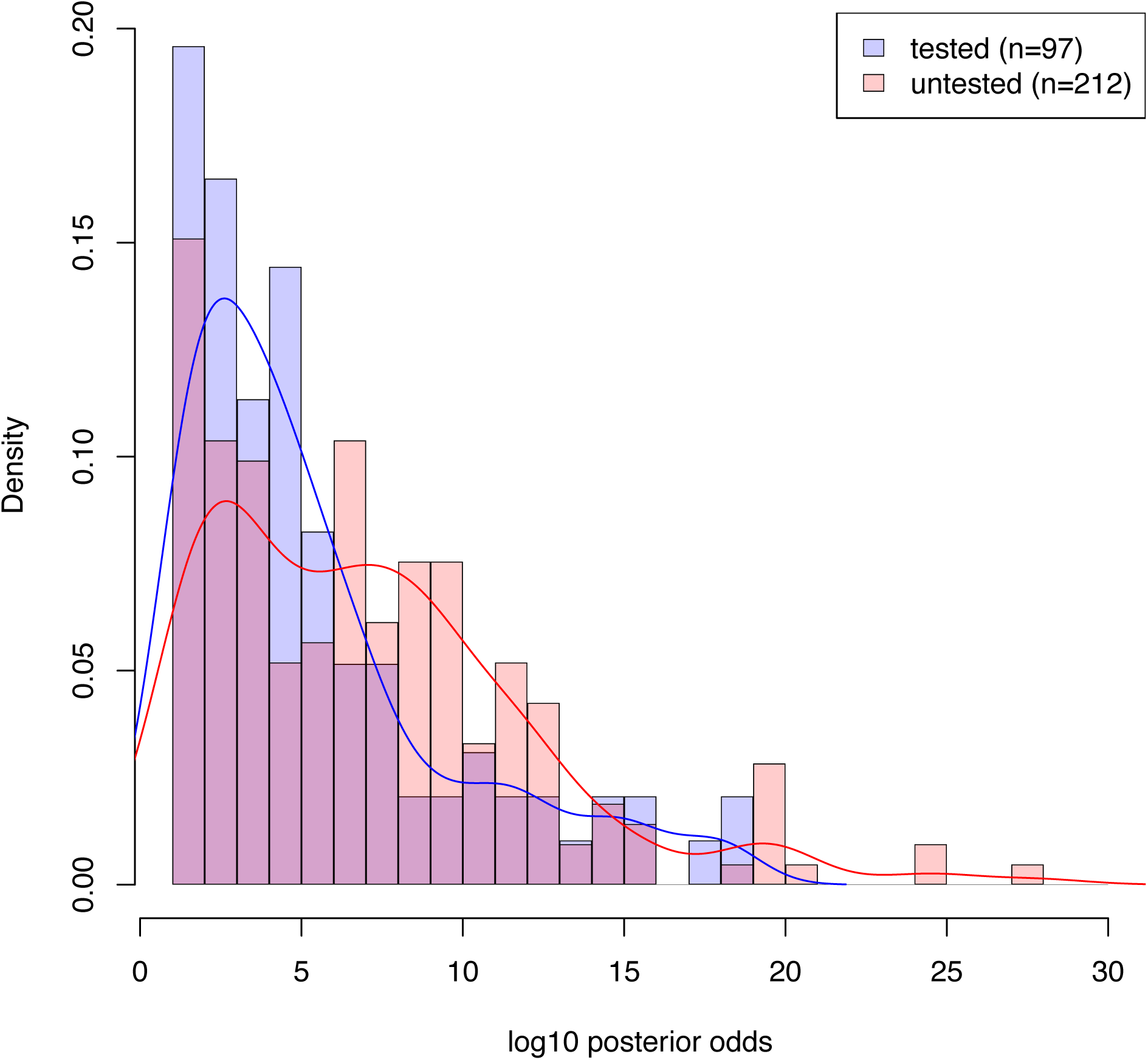
Posterior odds comparison for tested vs. untested mosaics. Among 309 candidates with EM-mosaic posterior odds scores available, we compared the distribution of tested (n=97) vs. untested (n=212) mosaics. The log_10_-scaled posterior odds distribution for the tested group is shown in blue (mean=5.382). The log_10_-scaled mean posterior odds for the untested group is shown in red (mean=7.050). The selected candidates had lower posterior odds than those not selected for confirmation (Mann Whitney *U* test *P*=0.002).

**Table S1. Proband Phenotypes**. Cardiac and neurodevelopmental phenotypes for CHD probands. NDD diagnosis is unknown for patients <1 year of age.

**Table S2. Cohort Summary**. Number of *de novo* and mosaic variants for probands with isolated CHD, extracardiac anomalies (ECA), neurodevelopmental delay (NDD) or unknown phenotypes.

**Table S3. EM-mosaic Mosaic Candidates**. Candidate mosaic variants identified by EM-mosaic and MiSeq validation results.

**Table S4. MosaicHunter Mosaic Candidates**. Candidate mosaic variants identified by MosaicHunter and MiSeq validation results.

**Table S5. Blood mosaic Candidate Validation by MiSeq.** 143 candidate mosaic sites were assessed using targeted PCR and deep sequencing. 85/97 (88%) of selected EM-mosaic sites and 44/68 (67%) of selected MosaicHunter sites were confirmed.

**Table S6. Cardiovascular Tissues with Whole Exome Sequencing**. 70 tissues from 66 CHD probands were assessed for mosaic variants.

**Table S7. CHD tissue mosaic Candidate Validation by MiSeq.** 24 candidate mosaic sites were assessed using targeted PCR and deep sequencing. 85/92 (92%) of selected EM-mosaic sites and 44/64 (69%) of selected MosaicHunter sites were confirmed .

**Table S8. CHD related genes.** We considered the union of genes highly expressed in the developing heart (HHE) and known candidate CHD genes {Jin 2017} as CHD-related genes (n=4558).

**Table S9. All Protein-Altering Mosaics**. Detailed information for 398 mosaic variants predicted to affect protein sequence.

**Table S10. Damaging Mosaics in CHD-Relevant Genes**. Detailed information for 25 mosaic variants likely to contribute to CHD.

## Declarations

### Ethics approval and consent to participate

CHD subjects were recruited to the Congenital Heart Disease Network Study of the Pediatric Cardiac Genomics Consortium (CHD GENES: ClinicalTrials.gov identifier <u>NCT01196182</u>). The institutional review boards of Boston’s Children’s Hospital, Brigham and Women’s Hospital, Great Ormond Street Hospital, Children’s Hospital of Los Angeles, Children’s Hospital of Philadelphia, Columbia University Medical Center, Icahn School of Medicine at Mount Sinai, Rochester School of Medicine and Dentistry, Steven and Alexandra Cohen Children’s Medical Center of New York, and Yale School of Medicine approved the protocols. All subjects or their parents provided informed consent.

### Consent for publication

See above.

### Availability of data and material

EM-mosaic and code for analyzing data are available from https://github.com/ShenLab/mosaicism. The MosaicHunter software is available from http://mosaichunter.cbi.pku.edu.cn/. SAMtools is available from http://www.htslib.org/. ANNOVAR is available from http://annovar.openbioinformatics.org/en/latest/. Integrative Genomics Viewer (IGV) software is available from https://software.broadinstitute.org/software/igv/. Whole-exome sequencing data have been deposited in the database of Genotypes and Phenotypes (dbGaP) under accession numbers <u>phs000571.v1.p1</u>, <u>phs000571.v2.p1</u> and <u>phs000571.v3.p2</u>. In-house pipelines are available from the corresponding authors on reasonable request.

### Authors’ contributions

YS, JGS, CES, and WKC conceived and oversaw the study. AH, SUM, JALW, HQ, KBM, JGS, CES, YS, WKC analyzed the data. AH developed the EM-mosaic pipeline and wrote the statistical analysis code. SUM, JALW carried out MosaicHunter analyses of blood and tissue samples. JMG, AT, SD performed MiSeq experimental confirmation. AH, SUM, EG, CES, WKC interpreted the impact of mosaics on participant clinical phenotypes. DB, RWK, JWN, GAP, DS, MT-F, MB, RPL, EG, BDG, CES, JGS, WKC were involved in cohort ascertainment, phenotypic characterization, and recruitment. DM collected cardiovascular tissue samples. AH, SUM, JALW, YS, JGS, CES, WKC wrote the manuscript. All authors read and approved the manuscript.

## Acknowledgements

The authors would like to thank all the participants and their families.

## References

Acuna-Hidalgo, R., Bo, T., Kwint, M. P., van de Vorst, M., Pinelli, M., Veltman, J. A., … Gilissen, C. (2015). Post-zygotic Point Mutations Are an Underrecognized Source of De Novo Genomic Variation. The American Journal of Human Genetics, 97(1), 67–74. http://doi.org/10.1016/J.AJHG.2015.05.008

Belickova, M., Vesela, J., Jonasova, A., Pejsova, B., Votavova, H., Merkerova, M. D., … Cermak, J. (2016). TP53 mutation variant allele frequency is a potential predictor for clinical outcome of patients with lower-risk myelodysplastic syndromes. Oncotarget, 7(24), 36266–36279. http://doi.org/10.18632/oncotarget.9200

Biesecker, L. G., & Spinner, N. B. (2013). A genomic view of mosaicism and human disease. Nature Reviews Genetics, 14(5), 307–320. http://doi.org/10.1038/nrg3424

Briggs, L. E., Kakarla, J., & Wessels, A. (2012). The pathogenesis of atrial and atrioventricular septal defects with special emphasis on the role of the dorsal mesenchymal protrusion. Differentiation, 84(1), 117–130. http://doi.org/10.1016/j.diff.2012.05.006

Cai, C.-L., Liang, X., Shi, Y., Chu, P.-H., Pfaff, S. L., Chen, J., & Evans, S. (2003). Isl1 identifies a cardiac progenitor population that proliferates prior to differentiation and contributes a majority of cells to the heart. Developmental Cell, 5(6), 877–89. Retrieved from http://www.ncbi.nlm.nih.gov/pubmed/14667410

Cibulskis, K., Lawrence, M. S., Carter, S. L., Sivachenko, A., Jaffe, D., Sougnez, C., … Getz, G. (2013). Sensitive detection of somatic point mutations in impure and heterogeneous cancer samples. Nature Biotechnology, 31(3), 213–219. http://doi.org/10.1038/nbt.2514

Cohn, D. H., Starman, B. J., Blumberg, B., & Byers, P. H. (1990). Recurrence of lethal osteogenesis imperfecta due to parental mosaicism for a dominant mutation in a human type I collagen gene (COL1A1). American Journal of Human Genetics, 46(3), 591–601. Retrieved from http://www.ncbi.nlm.nih.gov/pubmed/2309707

Colombo, S., de Sena-Tomás, C., George, V., Werdich, A. A., Kapur, S., MacRae, C. A., & Targoff, K. L. (2017). *nkx* genes establish SHF cardiomyocyte progenitors at the arterial pole and pattern the venous pole through Isl1 repression. Development, dev.161497. http://doi.org/10.1242/dev.161497

Dawson, K., Aflaki, M., & Nattel, S. (2013). Role of the Wnt-Frizzled system in cardiac pathophysiology: A rapidly developing, poorly understood area with enormous potential. Journal of Physiology, 591(6), 1409–1432. http://doi.org/10.1113/jphysiol.2012.235382

DePristo, M. A., Banks, E., Poplin, R., Garimella, K. V, Maguire, J. R., Hartl, C., … Daly, M. J. (2011). A framework for variation discovery and genotyping using next-generation DNA sequencing data. Nature Genetics, 43(5), 491–8. http://doi.org/10.1038/ng.806

Donkervoort, S., Hu, Y., Stojkovic, T., Voermans, N. C., Foley, A. R., Leach, M. E., … Bönnemann, C. G. (2015). Mosaicism for Dominant Collagen 6 Mutations as a Cause for Intrafamilial Phenotypic Variability. Human Mutation, 36(1), 48–56. http://doi.org/10.1002/humu.22691

Dou, Y., Yang, X., Li, Z., Wang, S., Zhang, Z., Ye, A. Y., … Wei, L. (2017). Postzygotic single-nucleotide mosaicisms contribute to the etiology of autism spectrum disorder and autistic traits and the origin of mutations. Human Mutation, 38(8), 1002–1013. http://doi.org/10.1002/humu.23255

Drake, K. M., Comhair, S. A., Erzurum, S. C., Tuder, R. M., & Aldred, M. A. (2015). Endothelial Chromosome 13 Deletion in Congenital Heart Disease–associated Pulmonary Arterial Hypertension Dysregulates SMAD9 Signaling. American Journal of Respiratory and Critical Care Medicine, 191(7), 850–854.

Etheridge, S. P., Bowles, N. E., Arrington, C. B., Pilcher, T., Rope, A., Wilde, A. A. M., … Tristani-Firouzi, M. (2011). Somatic mosaicism contributes to phenotypic variation in Timothy syndrome. American Journal of Medical Genetics Part A, 155(10), 2578–2583. http://doi.org/10.1002/ajmg.a.34223

Finger, J. H., Smith, C. M., Hayamizu, T. F., McCright, I. J., Xu, J., Law, M., … Ringwald, M. (2017). The mouse Gene Expression Database (GXD): 2017 update. Nucleic Acids Research, 45(D1), D730–D736. http://doi.org/10.1093/nar/gkw1073

Fischbach, G. D., & Lord, C. (2010). The Simons Simplex Collection: A Resource for Identification of Autism Genetic Risk Factors. Neuron, 68(2), 192–195. http://doi.org/10.1016/j.neuron.2010.10.006

Francioli, L. C., Polak, P. P., Koren, A., Menelaou, A., Chun, S., Renkens, I., … Sunyaev, S. R. (2015). Genome-wide patterns and properties of de novo mutations in humans. Nature Genetics, 47(7), 822–826. https://doi.org/10.1038/ng.3292

Freed, D., & Pevsner, J. (2016). The Contribution of Mosaic Variants to Autism Spectrum Disorder. PLOS Genetics, 12(9), e1006245. http://doi.org/10.1371/journal.pgen.1006245

Fryxell, K. J., & Moon, W.-J. (2005). CpG Mutation Rates in the Human Genome Are Highly Dependent on Local GC Content. Molecular Biology and Evolution, 22(3), 650–658. https://doi.org/10.1093/molbev/msi043

Genovese, G., Kähler, A. K., Handsaker, R. E., Lindberg, J., Rose, S. A., Bakhoum, S. F., … McCarroll, S. A. (2014). Clonal Hematopoiesis and Blood-Cancer Risk Inferred from Blood DNA Sequence. New England Journal of Medicine, 371(26), 2477–2487. http://doi.org/10.1056/NEJMoa1409405

Ghedira, N., Kraoua, L., Lagarde, A., Abdelaziz, R. Ben, Olschwang, S., Desvignes, J. P., … Mrad, R. (2017). Further Evidence for the Implication of LZTR1, a Gene not Associated with the Ras-Mapk Pathway, in the Pathogenesis of Noonan Syndrome. Biology and Medicine, 09(06), 4–7. http://doi.org/10.4172/0974-8369.1000414

Giampietro, C., Deflorian, G., Gallo, S., Di Matteo, A., Pradella, D., Bonomi, S., … Ghigna, C. (2015). The alternative splicing factor Nova2 regulates vascular development and lumen formation. Nature Communications, 6, 1–15. http://doi.org/10.1038/ncomms9479

Golzio, C., Havis, E., Daubas, P., Nuel, G., Babarit, C., Munnich, A., … Etchevers, H. C. (2012). ISL1 Directly Regulates FGF10 Transcription during Human Cardiac Outflow Formation. PLoS ONE, 7(1), e30677. http://doi.org/10.1371/journal.pone.0030677

Happle, R. (1986). The McCune-Albright syndrome: a lethal gene surviving by mosaicism. Clinical Genetics, 29(4), 321–4. Retrieved from http://www.ncbi.nlm.nih.gov/pubmed/3720010

Happle, R. (1993). Mosaicism in human skin. Understanding the patterns and mechanisms. Archives of Dermatology, 129(11), 1460–70. Retrieved from http://www.ncbi.nlm.nih.gov/pubmed/8239703

Heinrich, V., Stange, J., Dickhaus, T., Imkeller, P., Krüger, U., Bauer, S., … Krawitz, P. M. (2012). The allele distribution in next-generation sequencing data sets is accurately described as the result of a stochastic branching process. Nucleic Acids Research, 40(6), 2426–2431. http://doi.org/10.1093/nar/gkr1073

Homsy, J., Zaidi, S., Shen, Y., Ware, J. S., Samocha, K. E., Karczewski, K. J., … Chung, W. K. (2015). De novo mutations in congenital heart disease with neurodevelopmental and other congenital anomalies. Science, 350(6265), 1262–1266. http://doi.org/10.1126/science.aac9396

Hu, M., Sun, X.-J., Zhang, Y.-L., Kuang, Y., Hu, C.-Q., Wu, W.-L., … Chen, Z. (2010). Histone H3 lysine 36 methyltransferase Hypb/Setd2 is required for embryonic vascular remodeling. Proceedings of the National Academy of Sciences, 107(7), 2956–2961. http://doi.org/10.1073/pnas.0915033107

Huang, A. Y., Zhang, Z., Ye, A. Y., Dou, Y., Yan, L., Yang, X., … Wei, L. (2017). MosaicHunter: accurate detection of postzygotic single-nucleotide mosaicism through next-generation sequencing of unpaired, trio, and paired samples. Nucleic Acids Research, 45(10), e76–e76. http://doi.org/10.1093/nar/gkx024

Ioannidis, N. M., Rothstein, J. H., Pejaver, V., Middha, S., McDonnell, S. K., Baheti, S., … Sieh, W. (2016). REVEL: An Ensemble Method for Predicting the Pathogenicity of Rare Missense Variants. The American Journal of Human Genetics, 99(4), 877–885. http://doi.org/10.1016/j.ajhg.2016.08.016

Jaiswal, S., Fontanillas, P., Flannick, J., Manning, A., Grauman, P. V., Mar, B. G., … Ebert, B. L. (2014). Age-Related Clonal Hematopoiesis Associated with Adverse Outcomes. New England Journal of Medicine, 371(26), 2488–2498. http://doi.org/10.1056/NEJMoa1408617

Jamuar, S. S., Lam, A.-T. N., Kircher, M., D’Gama, A. M., Wang, J., Barry, B. J., … Walsh, C. A. (2014). Somatic Mutations in Cerebral Cortical Malformations. New England Journal of Medicine, 371(8), 733–743. http://doi.org/10.1056/NEJMoa1314432

Jin, S. C., Homsy, J., Zaidi, S., Lu, Q., Morton, S., DePalma, S. R., … Brueckner, M. (2017). Contribution of rare inherited and de novo variants in 2,871 congenital heart disease probands. Nature Genetics, 49(11), 1593–1601. http://doi.org/10.1038/ng.3970

Krupp, D. R., Barnard, R. A., Duffourd, Y., Evans, S. A., Mulqueen, R. M., Bernier, R., … O’Roak, B. J. (2017). Exonic Mosaic Mutations Contribute Risk for Autism Spectrum Disorder. The American Journal of Human Genetics, 101(3), 369–390. http://doi.org/10.1016/j.ajhg.2017.07.016

Kurahashi, H., Akagi, K., Inazawa, J., Ohta, T., Niikawa, N., Kayatani, F., … Nishisho, I. (1995). Isolation and characterization of a novel gene deleted in DiGeorge syndrome. Human Molecular Genetics, 4(4), 541–9. Retrieved from http://www.ncbi.nlm.nih.gov/pubmed/7633402

Kurek, K. C., Luks, V. L., Ayturk, U. M., Alomari, A. I., Fishman, S. J., Spencer, S. A., … Warman, M. L. (2012). Somatic Mosaic Activating Mutations in PIK3CA Cause CLOVES Syndrome. The American Journal of Human Genetics, 90(6), 1108–1115. http://doi.org/10.1016/j.ajhg.2012.05.006

Lauriat, T. L., Shiue, L., Haroutunian, V., Verbitsky, M., Ares, M., Ospina, L., & McInnes, L. A. (2008). Developmental expression profile ofquaking, a candidate gene for schizophrenia, and its target genes in human prefrontal cortex and hippocampus shows regional specificity. Journal of Neuroscience Research, 86(4), 785–796. http://doi.org/10.1002/jnr.21534

Lee, J. H., Huynh, M., Silhavy, J. L., Kim, S., Dixon-Salazar, T., Heiberg, A., … Gleeson, J. G. (2012). De novo somatic mutations in components of the PI3K-AKT3-mTOR pathway cause hemimegalencephaly. Nature Genetics, 44(8), 941–945. http://doi.org/10.1038/ng.2329

Lek, M., Karczewski, K. J., Minikel, E. V., Samocha, K. E., Banks, E., Fennell, T., … Consortium, E. A. (2016). Analysis of protein-coding genetic variation in 60,706 humans. Nature, 536(7616), 285–291. http://doi.org/10.1038/nature19057

Li, H., Handsaker, B., Wysoker, A., Fennell, T., Ruan, J., Homer, N., … 1000 Genome Project Data Processing Subgroup. (2009). The Sequence Alignment/Map format and SAMtools. Bioinformatics, 25(16), 2078–2079. http://doi.org/10.1093/bioinformatics/btp352

Lim, E. T., Uddin, M., De Rubeis, S., Chan, Y., Kamumbu, A. S., Zhang, X., … Walsh, C. A. (2017). Rates, distribution and implications of postzygotic mosaic mutations in autism spectrum disorder. Nature Neuroscience, 20(9), 1217–1224. http://doi.org/10.1038/nn.4598

Manheimer, K. B., Richter, F., Edelmann, L. J., D’Souza, S. L., Shi, L., Shen, Y., … Gelb, B. D. (2018). Robust identification of mosaic variants in congenital heart disease. Human Genetics, 137(2), 183–193. http://doi.org/10.1007/s00439-018-1871-6

McKenna, A., Hanna, M., Banks, E., Sivachenko, A., Cibulskis, K., Kernytsky, A., … DePristo, M. A. (2010). The Genome Analysis Toolkit: a MapReduce framework for analyzing next-generation DNA sequencing data. Genome Research, 20(9), 1297–303. http://doi.org/10.1101/gr.107524.110

Moorman, A., Webb, S., Brown, N. A., Lamers, W., & Anderson, R. H. (2003). Development of the heart: (1) formation of the cardiac chambers and arterial trunks. Heart (British Cardiac Society), 89(7), 806–14. Retrieved from http://www.ncbi.nlm.nih.gov/pubmed/12807866

Nawa, M., & Matsuoka, M. (2013). KCTD20, a relative of BTBD10, is a positive regulator of Akt. BMC Biochemistry, 14(1), 27. http://doi.org/10.1186/1471-2091-14-27

Noveroske, J. K., Lai, L., Gaussin, V., Northrop, J. L., Nakamura, H., Hirschi, K. K., & Justice, M. J. (2002). Quaking is essential for blood vessel development. Genesis (New York, N.Y.□: 2000), 32(3), 218–30. Retrieved from http://www.ncbi.nlm.nih.gov/pubmed/11892011

Poduri, A., Evrony, G. D., Cai, X., Elhosary, P. C., Beroukhim, R., Lehtinen, M. K., … Walsh, C. A. (2012). Somatic Activation of AKT3 Causes Hemispheric Developmental Brain Malformations. Neuron, 74(1), 41–48. http://doi.org/10.1016/j.neuron.2012.03.010

Ramsdell, A. F. (2005). Left–right asymmetry and congenital cardiac defects: Getting to the heart of the matter in vertebrate left–right axis determination. Developmental Biology, 288(1), 1–20. http://doi.org/10.1016/J.YDBIO.2005.07.038

Ramu, A., Noordam, M. J., Schwartz, R. S., Wuster, A., Hurles, M. E., Cartwright, R. A., & Conrad, D. F. (2013). DeNovoGear: de novo indel and point mutation discovery and phasing. Nature Methods, 10(10), 985–987. http://doi.org/10.1038/nmeth.2611

Ren, K., Yuan, J., Yang, M., Gao, X., Ding, X., Zhou, J., … Zhang, J. (2014). KCTD10 Is Involved in the Cardiovascular System and Notch Signaling during Early Embryonic Development. PLoS ONE, 9(11), e112275. http://doi.org/10.1371/journal.pone.0112275

Rivière, J.-B., Mirzaa, G. M., O’Roak, B. J., Beddaoui, M., Alcantara, D., Conway, R. L., … Dobyns, W. B. (2012). De novo germline and postzygotic mutations in AKT3, PIK3R2 and PIK3CA cause a spectrum of related megalencephaly syndromes. Nature Genetics, 44(8), 934–940. http://doi.org/10.1038/ng.2331

Sallman, D. A., Komrokji, R., Vaupel, C., Cluzeau, T., Geyer, S. M., McGraw, K. L., … Padron, E. (2016). Impact of TP53 mutation variant allele frequency on phenotype and outcomes in myelodysplastic syndromes. Leukemia, 30(3), 666–673. http://doi.org/10.1038/leu.2015.304

Sampson, J., Jacobs, K., Yeager, M., Chanock, S., & Chatterjee, N. (2011). Efficient study design for next generation sequencing. Genetic Epidemiology, 35(4), n/a-n/a. http://doi.org/10.1002/gepi.20575

Smith, C. L., Blake, J. A., Kadin, J. A., Richardson, J. E., Bult, C. J., & Mouse Genome Database Group. (2018). Mouse Genome Database (MGD)-2018: knowledgebase for the laboratory mouse. Nucleic Acids Research, 46(D1), D836–D842. http://doi.org/10.1093/nar/gkx1006

Smith, K. S., Yadav, V. K., Pei, S., Pollyea, D. A., Jordan, C. T., & De, S. (2016). SomVarIUS: somatic variant identification from unpaired tissue samples. Bioinformatics, 32(6), 808–813. http://doi.org/10.1093/bioinformatics/btv685

Soubrier, F., Chung, W. K., Machado, R., Grünig, E., Aldred, M., Geraci, M., … Humbert, M. (2013). Genetics and Genomics of Pulmonary Arterial Hypertension. Journal of the American College of Cardiology, 62(25), D13–D21. http://doi.org/10.1016/J.JACC.2013.10.035

Stevens, K. N., Hakonarson, H., Kim, C. E., Doevendans, P. A., Koeleman, B. P. C., Mital, S., … Gruber, P. J. (2010). Common Variation in ISL1 Confers Genetic Susceptibility for Human Congenital Heart Disease. PLoS ONE, 5(5), e10855. http://doi.org/10.1371/journal.pone.0010855

Stosser, M. B., Lindy, A. S., Butler, E., Retterer, K., Piccirillo-Stosser, C. M., Richard, G., & McKnight, D. A. (2018). High frequency of mosaic pathogenic variants in genes causing epilepsy-related neurodevelopmental disorders. Genetics in Medicine, 20(4), 403–410. http://doi.org/10.1038/gim.2017.114

Sun, J. X., He, Y., Sanford, E., Montesion, M., Frampton, G. M., Vignot, S., … Yelensky, R. (2018). A computational approach to distinguish somatic vs. germline origin of genomic alterations from deep sequencing of cancer specimens without a matched normal. PLOS Computational Biology, 14(2), e1005965. http://doi.org/10.1371/journal.pcbi.1005965

Tong, X., Zu, Y., Li, Z., Li, W., Ying, L., Yang, J., … Zhang, B. (2014). Kctd10 regulates heart morphogenesis by repressing the transcriptional activity of Tbx5a in zebrafish. Nature Communications, 5, 1–10. http://doi.org/10.1038/ncomms4153

Wallis, G. A., Starman, B. J., Zinn, A. B., & Byers, P. H. (1990). Variable expression of osteogenesis imperfecta in a nuclear family is explained by somatic mosaicism for a lethal point mutation in the alpha 1(I) gene (COL1A1) of type I collagen in a parent. American Journal of Human Genetics, 46(6), 1034–40. Retrieved from http://www.ncbi.nlm.nih.gov/pubmed/2339700

Weinstein, M. M., Kang, T., Lachman, R. S., Bamshad, M., Nickerson, D. A., Krakow, D., & Cohn, D. H. (2016). Somatic mosaicism for a lethal *TRPV4* mutation results in non-lethal metatropic dysplasia. American Journal of Medical Genetics Part A, 170(12), 3298–3302. http://doi.org/10.1002/ajmg.a.37942

Yamamoto, G. L., Aguena, M., Gos, M., Hung, C., Pilch, J., Fahiminiya, S., … Bertola, D. R. (2015). Rare variants in SOS2 and LZTR1 are associated with Noonan syndrome. Journal of Medical Genetics, 52(6), 413–421. http://doi.org/10.1136/jmedgenet-2015-103018

Zaidi, S., & Brueckner, M. (2017). Genetics and Genomics of Congenital Heart Disease. Circulation Research, 120(6), 923–940. http://doi.org/10.1161/CIRCRESAHA.116.309140

Zaidi, S., Choi, M., Wakimoto, H., Ma, L., Jiang, J., Overton, J. D., … Lifton, R. P. (2013). De novo mutations in histone-modifying genes in congenital heart disease. Nature, 498(7453), 220–223. http://doi.org/10.1038/nature12141

